# Museum genomics illuminate the high specificity of a bioluminescent symbiosis across a genus of reef fish

**DOI:** 10.1101/2020.10.01.323204

**Authors:** Alison L. Gould, Allison Fritts-Penniman, Ana Gaisiner

## Abstract

Symbiotic relationships between bioluminescent bacteria and fishes have evolved multiple times across hundreds of fish taxa, but relatively little is known about the specificity of these associations and how conserved they have been through time. This study describes the degree of specificity of a bioluminescent symbiosis between cardinalfishes in the genus *Siphamia* and luminous bacteria in the Vibrio family. Primarily using museum specimens, we investigate the co-divergence of host and symbiont and test for patterns of divergence that correlate with both biogeography and time. Contrary to expectations, we determined that the light organ symbionts of all 14 *Siphamia* species examined belong to one genetic clade of *Photobacterium mandapamensis* (Clade II), indicating that the association is highly specific and conserved across the host genus. Thus, we did not find evidence of codivergence among hosts and symbionts. We did observe that symbionts hosted by individuals sampled from colder water regions were more divergent, containing more than three times as many single nucleotide polymorphisms than the rest of the symbionts examined. Overall our findings indicate that the symbiosis between *Siphamia* fishes and *P. mandapamensis* Clade II has been highly conserved across a broad geographic range and through time despite the facultative nature of the bacterial symbiont. These results suggest that this bioluminescent symbiosis could have played a key role in the evolution of the host genus and that there are conserved mechanisms regulating its specificity that have yet to be defined.

## Introduction

Environmentally transmitted microbial symbionts are acquired by a host from a genetically diverse, free-living population of bacteria. These facultative symbionts must retain the genetic machinery necessary to associate with their hosts, while also being able to compete with the rest of the microbial community in the surrounding environment (Bright and Bulgheresi 2010). In the marine environment abiotic factors such as water flow and temperature play critically important roles in structuring the microbial community (Galand et al., 2010, Brown et al., 2012), and accordingly, the available free-living symbiont pool. Nevertheless, a colossal diversity of bacteria remains available to marine hosts, yet the associations between hosts and their microbial symbionts are highly specific; much more so than what would be expected based solely on diversity in the surrounding seawater (Trousselier et al., 2017). Thus, the combined influence of host attributes and abiotic factors contributes to the complexity of the specificity of environmentally transmitted host-symbiont associations, providing the opportunity to study the evolution of specificity and co-diversification of these critical associations.

Bioluminescent symbioses have evolved multiple times across diverse squid and fish taxa, including at least 17 times in the ray-finned fishes (Davis et al., 2016; Dunlap and Urbanczyk, 2013). Approximately 500 species of fish are known to be symbiotically bioluminescent, but our understanding of specificity between fish hosts and their bacterial symbionts is just emerging. Existing evidence suggests that some level of specificity between a host and luminous symbiont is maintained, at least at the host family level. For example, leiognathid fishes exclusively host *Photobacterium leiognathi* and *P. mandapamensis* (Kaeding et al. 2007), and ceratioid anglerfishes, representing four different host families sampled over a broad geographic range, hosted only two bacterial species, *Enterovibrio escacola* and *E. luxaltus* (Baker et al. 2019). Specificity has been described in 35 additional fish host species, comprising 7 families (Dunlap et al. 2007). This host family level of bacterial specificity is believed to result from the fish selecting for its particular symbiont while also preventing other bacteria from colonizing its light organ (Reichelt et al., 1977). Although fish hosts only associate with a narrow range of luminous bacteria, the symbionts are generally not obligately dependent on their host (but see Hendry et al., 2014) and can survive in a variety of other habitats including seawater, sediment, and the surfaces and digestive tracts of various marine organisms. Thus, the specificity of bioluminescent symbioses depends largely on host selectivity and the genetics of the association.

Within the cardinalfish family (Perciformes: Apogonidae), bioluminescence has evolved multiple times, however only species in the genus *Siphamia* rely on a symbiotic relationship with luminous bacteria to produce light; all other bioluminescent cardinalfishes produce their own light presumably via the acquisition of luciferin from their diet (Thacker and Roje 2009). All 25 species of *Siphamia* are symbiotically bioluminescent (Thacker and Roje 2009; Gon and Allen 2012). The fish possess a ventral light organ connected to the intestine, which functions to host a dense population of luminous bacteria (∼10^8^ cells) (Fig. 1) (Dunlap and Nakamura 2011). The symbionts are ingested by the host during larval development and subsequently colonize the host’s light organ (Dunlap et al., 2012).

**Figure 1.**
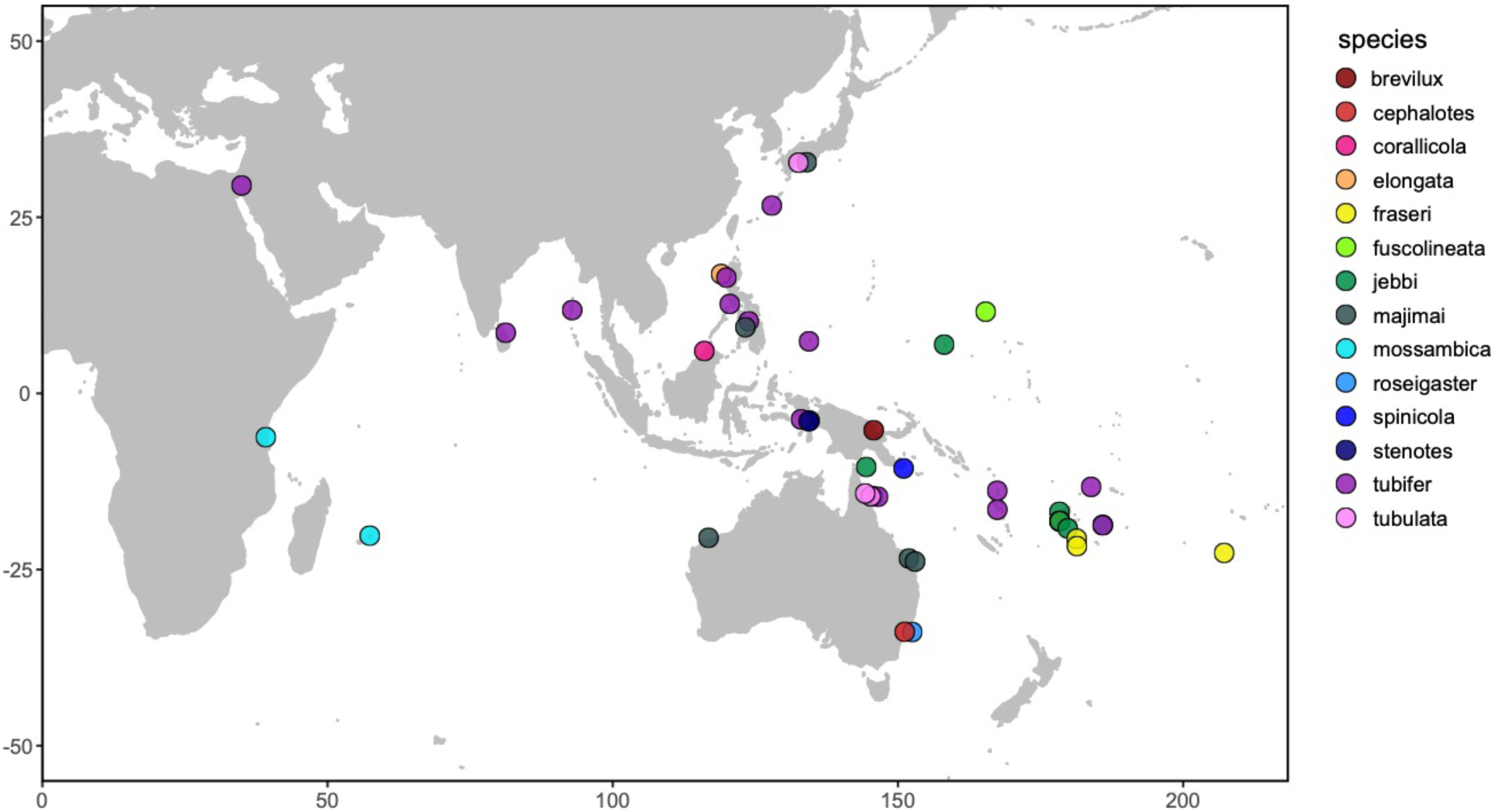
Map depicting the sampling locations of the Siphamia specimens examined in this study. Colors represent different *Siphamia* species as indicated in the figure legend.

The *Siphamia-Photobacterium* symbiosis readily lends itself to study both in the field and in the laboratory because, unlike most bioluminescent fish which occur in deep or open water environments, *Siphamia* reside in shallow waters with high habitat fidelity (Gould et al 2014). Furthermore, both host and symbiont can be readily cultured in captivity, making them ideal study organisms for laboratory investigations (Dunlap et al., 2012). However, the luminous symbionts of only one *Siphamia* species, S. *tubifer,* originating from a small geographic region in the Okinawa Islands, Japan, have been characterized to date; the specimens examined were found to host only members of Clade II of *Photobacterium mandapamensis* in their light organs, suggesting a high degree of specificity for this association (Kaeding et al., 2007, Gould and Dunlap 2019). *Siphamia tubifer* is broadly distributed throughout the Indo-Pacific, spanning from eastern Africa to the French Polynesian Islands (Gon and Allen 2012), thus the true degree of specificity across the geographic range of this association remains unknown. Furthermore, the luminous symbionts of the other 24 species in the host genus have yet to be identified.

The primary goals of this study were to characterize the degree of specificity of the bioluminescent symbiosis throughout the *Siphamia* genus and across the broad geographic range of S. *tubifer.* Taking advantage of previous collection efforts, we sampled geographically and temporally diverse *Siphamia* specimens (Fig. 2) from several natural history museums.

**Figure 2.**
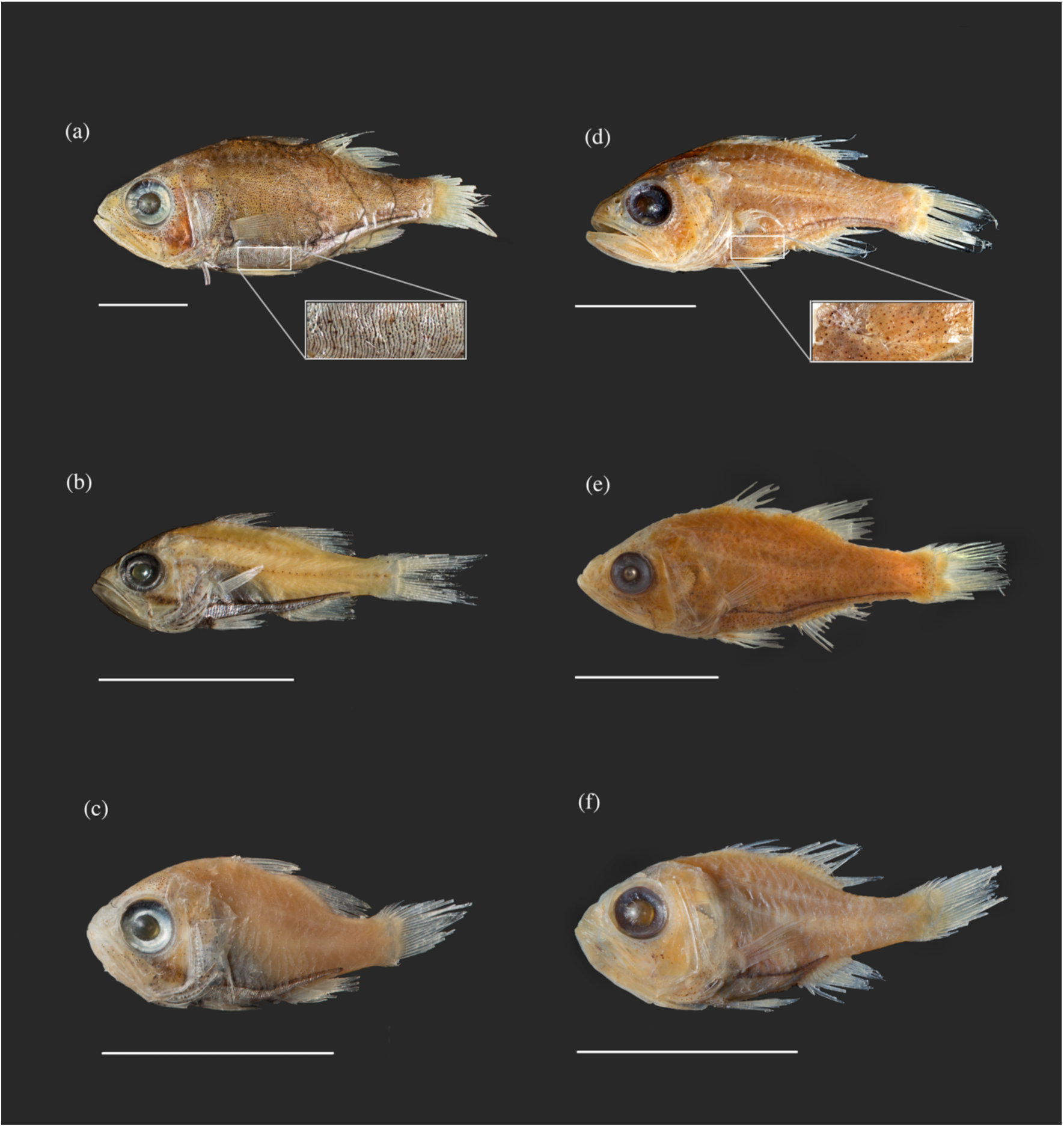
Photographs of select *Siphamia* specimens from lots used in this study. Specimens a-c represent the *tubifer* subgroup (Gon and Allen 2012), identified by the striated light organ (a) and specimens d-f represent the *tubulata* subgroup, identified by the spotted light organ (d). (a) S. *tubifer* (USNM341595) with insert of light organ detail showing striated morphology. (b) S. *stenotes* (USNM396981, paratype). (c) S. *jebbi* (CAS223855). (d) S. *tubulata* (CAS28515) with insert of light organ detail showing spotted morphology. (e) S. *corallicola* (USNM203781). (f) S. *brevilux* (CAS65338, paratype). Scale bars indicate 1 cm in length.

Recovering genetic information from wet specimens, particularly those initially fixed in formalin, is a new frontier in museum genomics. Here we present methods for extracting and sequencing the DNA of both a bacterial symbiont and its vertebrate host. Thus, we were able to test for evidence of co-diversification of host and its symbiont, and for patterns of symbiont diversity at the clade-level that correlate with biogeography, temperature, and time.

## Methods

### Taxon sampling and DNA extraction

We sampled 59 specimens representing 14 *Siphamia* species obtained from the combined wet collections of the California Academy of Sciences, the Australian Museum, and the Smithsonian National Museum of Natural History (Figs. 1–2, Table 1). To extract DNA from these specimens, we adapted the following protocol from two previous methods designed for use with formalin-fixed tissues (Ruane and Austin 2017, Hykin et al., 2015). Light organs were aseptically dissected and individually placed into 1 ml of GTE buffer and allowed to soak for three hours at room temperature. This step was repeated two times after which each light organ was transferred into a final 1 ml aliquot of fresh GTE buffer and left to soak overnight at room temperature. The following morning, each sample was transferred into 1 ml of 100% ethanol for one minute, followed by 1 ml of 70% ethanol for 5 minutes, and 1 ml of nuclease-free water for 10 minutes at room temperature. Light organs were then transferred into 180 ul of pre-heated (98°C) ATL buffer (QIAGEN) and incubated at 98°C for 15 minutes, after which samples were immediately placed on ice for at least 2 minutes. Once cooled, 40 ul of proteinase K was added to each sample and the samples were incubated at 60°C for 48 hours on a shaking heat block. Samples were vortexed periodically and additional 20 ul aliquots of proteinase K were added as needed (up to 100 ul total). Following this incubation period, DNA was extracted using the QIAGEN DNEasy Blood and Tissue Kit as described by the manufacturer. Purified DNA products were eluted into 50 ul of nuclease-free water after a 3-minute incubation at 55°C.

**Table 1.** Information for the *Siphamia* specimens sampled in this study. Listed are each specimen’s catalog number or unique identifier, species identification, sampling location and year, the minimum and maximum temperatures at that location, and the standard length of the individual sampled. Specimens with decimals after their catalog number or unique identifier indicate that more than one individual was sampled from the same specimen lot. Sea surface temperatures from the topmost meter of water at the geographical point of specimen collection were calculated as the temporal minimum and maximum from monthly climatologies (2002-2009) extracted from the Aqua-MODIS database available on Bio-ORACLE. (Tyberghein et al., 2012)

| Specimen ID | Species | Location | Year | Min Temp | Max Temp | Length (cm) |
| --- | --- | --- | --- | --- | --- | --- |
| AMI18353-041 | jebbi | Fiji | 1974 | 31.06 | 25.57 | 1.69 |
| AMI18740-066 | jebbi | Australia | 1975 | 29.48 | 24.69 | 1.46 |
| AMI19450-018.1 | tubifer | Australia | 1975 | 29.94 | 24.09 | 2.94 |
| AMI19450-018.2 | tubifer | Australia | 1975 | 29.94 | 24.09 | 3.62 |
| AMI20353-001 | majimai | Australia | 1972 | 31.76 | 22.61 | 1.67 |
| AMI20753-031 | tubulata | Australia | 1979 | 30.04 | 23.54 | 2.56 |
| AMI33715-016 | jebbi | Australia | 1993 | 29.70 | 25.08 | 1.46 |
| AMI37933-007 | tubifer | Vanuatu | 1997 | 30.18 | 27.30 | 2.19 |
| AMI40838-008 | cephalotes | Australia | 2001 | 23.03 | 15.49 | 3.07 |
| AMI40865-004.1 | roseigaster | Australia | 2001 | 24.16 | 18.64 | 4.56 |
| AMI40865-004.2 | roseigaster | Australia | 2001 | 24.16 | 18.64 | 4.55 |
| AMIB4208 | majimai | Australia | 1958 | 28.11 | 21.64 | 2.31 |
| AMIB4247 | tubifer | Vanuatu | 1959 | 29.64 | 26.23 | 2.01 |
| CAS247233.1 | mossambica | Zanzibar | 2018 | 31.06 | 25.92 | 2.55 |
| CAS247233.2 | mossambica | Zanzibar | 2018 | 31.06 | 25.92 | 2.92 |
| CAS222309 | jebbi | Fiji | 2002 | 30.28 | 26.16 | - |
| CAS223855 | jebbi | Fiji | 2002 | 29.88 | 25.89 | - |
| CAS223939.1 | jebbi | Fiji | 2002 | 29.88 | 25.89 | 2.35 |
| CAS223939.2 | jebbi | Fiji | 2002 | 29.88 | 25.89 | 1.81 |
| CAS223978.1 | unknown | Fiji | 2002 | 29.88 | 25.89 | 3.68 |
| CAS223978.2 | unknown | Fiji | 2002 | 29.88 | 25.89 | 4.05 |
| CAS223979.1 | fraseri | Fiji | 2002 | 29.88 | 25.89 | 2.8 |
| CAS223979.2 | fraseri | Fiji | 2002 | 29.88 | 25.89 | 3.04 |
| CAS225045 | jebbi | Fiji | 1999 | 29.88 | 25.89 | - |
| CAS27441 | tubifer | Philippines | 1931 | 30.77 | 27.89 | 3.26 |
| CAS28515 | tubulata | Australia | 1973 | 30.31 | 23.60 | - |
| CAS84356 | tubifer | Palau | 2012 | 30.33 | 28.57 | 1.9 |
| Stubifer_M118 | tubifer | Ryukyu Islands | 2013 | 29.60 | 20.83 | 1.3 |
| Smajimai_PVD | majimai | Japan | 2007 | 28.75 | 18.68 | 2.61 |
| Stubulata_PVD | tubulata | Japan | 2007 | 28.28 | 18.02 | 2.12 |
| Stubifer_S27 | tubifer | Ryukyu Islands | 2013 | 29.60 | 20.83 | 2.65 |
| Sstenotes_GRA.1 | stenotes | Indonesia | 2006 | 30.80 | 26.42 | 1.89 |
| Sstenotes_GRA.2 | stenotes | Indonesia | 2006 | 30.80 | 26.42 | 1.98 |
| Stubifer_GRA.1 | tubifer | Indonesia | 2006 | 30.88 | 26.60 | 2.39 |
| Stubifer_GRA.2 | tubifer | Indonesia | 2006 | 30.88 | 26.60 | 2.85 |
| USNM112099 | elongata | Philippines | 1909 | 30.74 | 27.58 | 3.46 |
| USNM142281.1 | fuscolineata | Marshall Islands | 1946 | 29.70 | 27.05 | 2.2 |
| USNM142281.2 | fuscolineata | Marshall Islands | 1946 | 29.70 | 27.05 | 2.76 |
| USNM203781 | corallicola | Borneo | 1965 | 31.33 | 28.34 | 2.58 |
| USNM223216 | jebbi | Micronesia | 1980 | 30.73 | 28.31 | 1.74 |
| USNM245638 | jebbi | Fiji | 1982 | 29.10 | 25.04 | 2.07 |
| USNM245641 | fraseri | Fiji | 1982 | 28.60 | 24.12 | 4.13 |
| USNM245642 | fraseri | Fiji | 1982 | 28.08 | 23.32 | 3.65 |
| USNM298542 | brevilux | Papua New Guinea | 1988 | 30.83 | 28.73 | 2.24 |
| USNM341594 | jebbi | Tonga | 1993 | 29.02 | 25.28 | 1.91 |
| USNM341595 | tubifer | Tonga | 1993 | 29.59 | 25.63 | 3.87 |
| USNM349778 | mossambica | Mauritius | 1995 | 28.76 | 23.68 | 2.36 |
| USNM357884 | tubifer | Philippines | 1980 | 30.60 | 27.53 | 3.68 |
| USNM357889 | spinicola | Papua New Guinea | 1975 | 29.83 | 25.76 | 3.11 |
| USNM357892 | tubifer | Red Sea | 1969 | 28.10 | 21.47 | 3.35 |
| USNM357897 | tubifer | Andaman | 1963 | 31.15 | 27.89 | 4.09 |
| USNM357999 | tubifer | Sri Lanka | 1970 | 30.58 | 27.33 | 2.94 |
| USNM358001 | majimai | Philippines | 1978 | 30.06 | 27.19 | 2.1 |
| USNM374480 | majimai | Australia | 1966 | 27.85 | 21.93 | 1.97 |
| USNM374837 | unknown | Wallis and Futuna | 2000 | 30.27 | 28.26 | 1.96 |
| USNM396981 | stenotes | Indonesia | 2006 | 31.57 | 27.68 | 1.89 |
| USNM412731 | jebbi | Philippines | 2003 | 30.79 | 28.54 | 1.73 |
| USNM430718 | fraseri | French Polynesia | 2013 | 27.80 | 23.49 | 3.34 |

### Library preparation and sequencing

Samples were quantified using the Qubit dsDNA HS Assay Kit on the Qubit 2.0 Fluorometer (Invitrogen) and profiled with an Agilent 2100 Bioanalyzer. Samples with a peak in size distribution greater than 300 bp were sonicated with a Qsonica (Q800R3) for one or two minutes (if peak was greater than 1,500 bp) with a pulse rate of 10-10 seconds and an amplitude of 25%. Samples were then treated with the NEBNext© FFPE DNA Repair Mix following the manufacturer’s instructions and DNA libraries were immediately prepped using the NEBNext© Ultra II DNA Library Prep Kit. Samples with low or undetectable quantities of dsDNA were re­quantified using the Qubit ssDNA HS Assay Kit and prepared using the Accel-NGS 1S Plus

DNA Library Kit (Swift Biosciences), which uses both single- and double-stranded DNA as templates. Each sample was uniquely indexed with the NEBNext© Multiplex Oligos for Illumina. Final libraries were cleaned with AMPure XP magnetic beads, pooled, and sequenced as single-end 150 bp (UC Berkeley, QB3) or paired-end 150 bp (NovoGene) reads on the Illumina HiSeq 4000 platform, or as paired-end 150 bp reads on the Illumina NovaSeq S4 platform (Genewiz). Table S1 contains details for each sample and library preparation.

### Sequence analysis

Sequences were demultiplexed, trimmed and quality filtered for a Phred score of 20 or above using Trimmomatic (Bolger et al., 2014). The remaining reads were aligned to the reference genome of *Photobacterium mandapamensis,* isolated from the light organ of *Siphamia tubifer* (Urbanczyk et al., 2011) with BWA-MEM (Li 2013). Unaligned sequences were then processed with MitoFinder (Allio et al., 2020) using the reference mitochondrial genome of the Banggai cardinalfish *Pterapogon kauderni* (Matias and Hereward 2018). All cardinalfish cytochrome oxidase subunit 1 *(COI)* gene sequences that were recovered aligned using MUSCLE (Edgar 2004) and a maximum likelihood analysis was carried out with raxml-ng (Kozlovet *al.,* 2019) using the evolutionary model TIM2+F+I+G4 which had the lowest BIC score as predicted by IQtree (Nguyen et al., 2015) and 1,000 bootstraps to infer the phylogenetic relationships between host species. *COI* sequences of *Siphamia* spp. from previous studies were also included in the analysis (Table S2). An additional phylogeny was inferred from a supermatrix of 15 mitochondrial genes *(ATP6, ATP8, COXI, COX2, COX3, CYTB, ND1, ND2, ND3, ND4, ND4L, ND5, ND6, 16S, 18S)* identified by MitoFInder that were present in at least 70% of the individuals included in the analysis using the SuperCRUNCH python toolkit (Portik and Wiens 2020). The concatenated supermatrix alignment was used in a maximum likelihood analysis by raxml-ng with 500 bootstrap replicates and the evolutionary model TIM2+F+R4 as predicted by IQtree to infer the phylogenetic relationships between species.

Two approaches were used to determine the identity of the light organ symbionts. First, 16S rRNA gene sequences were extracted from each data set by aligning all light organ sequences to the complete 16S sequence of a free-living strain of *Photobacterium leiognathi* (AY292917)(Nishiguchi and Nair 2003) with BWA-MEM (Li 2013). A sequence similarity search was then performed with the basic local alignment search tool (BLAST) (Altschul et al., 1990) against NCBI’s microbial database to identify the known sequence with the lowest E-value and highest percent identity. Second, the average nucleotide identity (ANI) of each sample was calculated relative to several *Photobacterium* species for which entire genome sequences are available from the NCBI genome database *(P. kishintanii* pjapo1.1 - NZ_PYNK00000000; *P. leiognathi* lrivu4.1 - NZ_BANQ00000000; *P. mandapamensis* ajapo4.1 - NZ_PYNQ01000000; *P. mandapamensis* gjord1.1 - NZ_PYNP00000000; *P. mandapamensis* svers1.1 - NZ_PYNT00000000) with the program fastANI (Jain et al., 2018).

To infer the phylogenetic relationships between symbionts from different hosts, all sequences that aligned to the reference genome of *P. mandapamensis* (Urbanczyk et al., 2011) were also analyzed for sequence variation between with the program snippy (Seemann 2015), requiring a minimum depth of 10x and a minimum percent of reads to be 90% to call a variant. A sequence alignment based on a core set of single nucleotide polymorphisms (SNPs) was then created across symbionts with enough genome coverage to produce a core set of at least 1,000 SNPs and including two additional reference genomes of *P. mandapamensis* representing both Clade I (ajapo4.1) and Clade II (Res4.1). The phylogenetic relationships of these bacteria were then inferred with raxml-ng (Kozlov et al., 2019) using the evolutionary model model TVM+F+R3, which had the lowest *BIC* score as predicted by IQtree (Nguyen et al., 2015) and 1,500 bootstrap replicates.

Samples included in both the host and symbiont phylogenies were then compared and tested for co-divergence using the cospeciation function in the R phytools package (Revell 2012). SNPs were annotated with the program SNPeff (Cingolani et al., 2012). Pairwise phylogenetic (patristic) distances between symbionts were calculated with the adephylo package (Jombart and Dray 2008) in R, and pairwise geographic distances were calculated based on each specimen’s latitude and longitude using the R package geodist (Padgham and Summer 2020). Tests for correlations between the phylogenetic distances for each pair of symbionts and their geographic distance or difference between sampling years were carried out, and P values were adjusted for multiple comparisons with the Holm method in R.

## Results

### DNA recovery

Variable amounts of total DNA were recovered from the light organs of preserved *Siphamia* specimens, ranging from undetectable levels (<2 ng) to more than 1,500 ng, and there was no correlation between DNA yield and specimen size (Spearman’s rank correlation: rho=0.52, P=0.09). Despite this variability in yield, quality DNA sequences were recovered from several specimens with undetectable levels of starting DNA. In fact, some samples with undetectable levels of input DNA resulted in >90% coverage of the symbiont genome at 10x depth. Of note, many of those sequence libraries were prepared using the Swift Bioscience Accel-NGS 1S Plus DNA Library Kit which uses both double and single stranded DNA as starting template (Table S1).

### Host phylogeny

Host *COI* sequences were recovered from 32 samples and analyzed with an additional 12 *Siphamia COI* sequences from previous studies (Table S2) to generate a maximum likelihood phylogeny of 17 *Siphamia* species (Fig 3). The supermatrix of 15 mitochondrial genes from 27 *Siphamia* specimens representing 11 species resulted in a phylogenetic tree with similar, but not identical, topology and stronger bootstrap support at the nodes (Figure S1).

**Figure 3.**
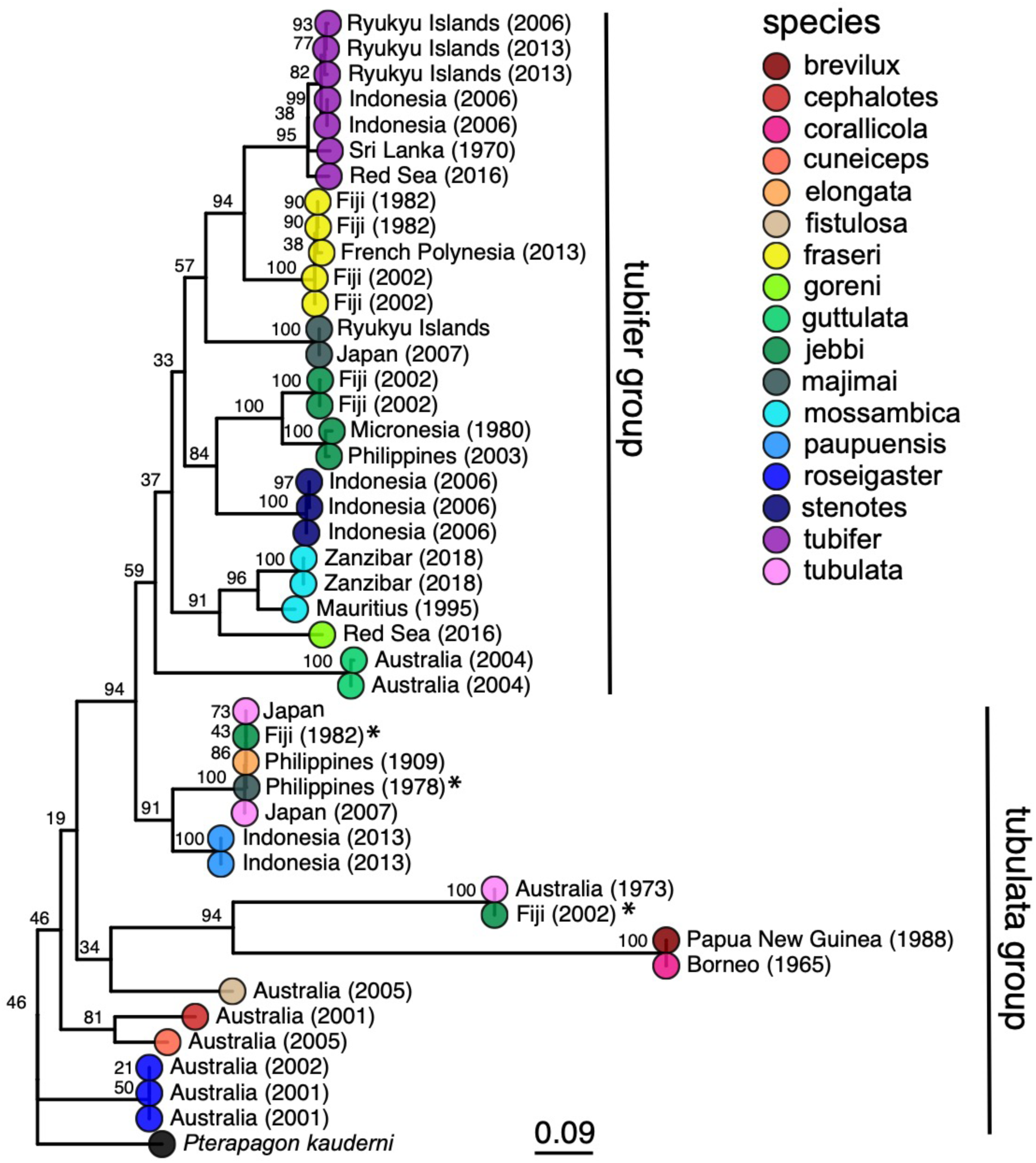
Maximum likelihood phylogeny of *Siphamia* based on *COI* gene sequences. Species identities are indicated by the branch tip colors and the sampling location and year of each specimen is listed in the branch label. Bootstrap support values are indicated at each node. The Banggai cardinalfish, *Pterapogon kauderni,* was used as the outgroup. The *tubifer* and *tubulata* subgroups within *Siphamia* (Gon & Allen 2012) are highlighted with vertical lines to the right of the tree. Specimens that fall outside of their designated subgroup based on species identities are indicated with an *.

Our phylogenetic hypothesis for *Siphamia* is very similar to that proposed by Gon and Allen (2012) using morphological characters, with slight variations in the placement of specific taxa. Our tree contains a clade that corresponds to Gon and Allen’s S. *tubifer* species group, characterized by a striated pattern on the light organ (Fig 2), although 1 individual S. *majimai* and 2 S. *jebbi* specimens fell out of this group (Fig 3). Within this group, our trees support the relationships of S. *jebbi* and S. *stenotes* as sister species, as well as S. *tubifer* and S. *fraseri.* The relative placement of S. *mossambica, S. majimai,* and S. *goreni* varies among the trees, but there is support for S. *mossambica* and S. *goreni* as sister species in the *COI* tree, S. *mossambica* and S. *majimai* as sisters in the supermatrix tree, and S. *majimai* and S. *goreni* as sisters in the morphological tree (Gon and Allen 2012). As such, it is likely that all of three of these species belong to one clade. The relationships among the species outside of the S. *tubifer* group are less certain, with several species clustering into species complexes. However, S. *roseigaster, S. cuneiceps,* and S. *cephalotes* consistently fall out near the base of the tree, indicating that these species diverged earlier.

### Symbiont identification and phylogeny

To identify the light organ symbionts of the *Siphamia* hosts, we recovered 16S rRNA sequences from the shotgun sequence data. 93% of all samples had >95% coverage of the 16S rRNA gene at 10x read depth. Samples were putatively identified by matching sequences against those in the NCBI database and 67% had *P. mandapamensis* as their top hit (Table S3.). All other symbionts were identified as *P. leiognathi.* However, previous analyses of 16S rRNA gene sequences could not resolve *P. leiognathi* from *P. mandapamensis* (Ast and Dunlap 2004, Wada *et al.,* 2006). Therefore, to confirm the identities of the light organ symbionts, we also calculated the average nucleotide identity (ANI) of each symbiont relative to several *Photobacterium* strains for which whole genomes are available. 86% of the symbionts examined had ANI values of 95% or greater relative to *P. mandapamensis* strains in Clade II (gjord 1.1 and svers1.1) (Table 2), which is the recommended value to delimit bacterial species (Goris et al., 2007). All remaining samples also had the highest ANI values relative to *P. mandapamensis* Clade II, with the exception of one sample (AMI18740-066), which was most similar to *P. mandapamensis* Clade I, however many of these samples also had low genome coverage (Table S1). None of the symbionts had ANI values relative to *P. leiognathi* that were higher than those relative to *P. mandapamensis.*

**Table 2.** Average nucleotide identities (%) of the light organ symbionts of the *Siphamia* specimens sampled in this study relative to several *Photobacterium* species for which entire genomes are available from NCBI: *P. kishitanii* (pjapo1.1), *P. leiognathi* (lrivu4.1), *P. mandapamensis,* Clade I (ajapo4.1), *P. mandapamensis,* Clade II (gjord 1.1, svers1.1). Values in bold are 94.5% or greater. Also listed is each specimen’s catalog number or unique identifier and the symbiont’s percent genome coverage at 10x sequencing depth relative to *P. mandapamensis* (svers1.1)

| Specimen ID | *pjapo1.1* | *lrvu4.1* | *ajapo4.1* | *gjord1.1* | *svers1.1* | %10x |
| --- | --- | --- | --- | --- | --- | --- |
| Stubulata_PVD | 80.1 | 92.9 | **96.7** | **97.4** | **97.4** | 95.1 |
| Smajimai_PVD | 80.1 | 92.8 | **96.7** | **97.4** | **97.3** | 94.8 |
| AMI40838-008 | 80.2 | 92.8 | **96.5** | **97.2** | **97.1** | 92.3 |
| AMI40865-004.1 | 80.1 | 92.9 | **96.5** | **97.1** | **97.2** | 93.4 |
| Stubifer_GRA.2 | 79.8 | 92.2 | **96** | **96.8** | **97.2** | 52 |
| CAS84356 | 80 | 92.1 | **96** | **96.6** | **97.1** | 64.8 |
| CAS247233.1 | 80 | 92 | **95.8** | **96.6** | **97** | 76.9 |
| Stubifer_S27 | 80 | 92.1 | **95.7** | **96.5** | **96.9** | 88.9 |
| USNM412731 | 80.2 | 91.9 | **95.6** | **96.3** | **96.4** | 95.9 |
| CAS27441 | 80 | 91.8 | **95.5** | **96.2** | **96.6** | 95.3 |
| AMI40865-004.2 | 79.7 | 92 | **95.7** | **96** | **96.2** | 5.9 |
| Sstenotes_GRA.1 | 80.1 | 91.8 | **95.4** | **96.1** | **96.2** | 95.8 |
| Sstenotes_GRA.2 | 79.6 | 91.6 | **95.2** | **95.9** | **96.5** | 0.5 |
| USNM245638 | 79.7 | 91.4 | **95.3** | **96** | **95.9** | 35.6 |
| USNM245641 | 79.9 | 91.4 | **95.1** | **95.7** | **95.9** | 97.5 |
| USNM245642 | 79.9 | 91.4 | **95.1** | **95.7** | **95.8** | 96.2 |
| CAS222309 | 79.9 | 91.3 | **95** | **95.7** | **95.8** | 95.2 |
| USNM223216 | 80 | 91.3 | **94.8** | **95.5** | **95.6** | 96.5 |
| USNM142281 | 79.7 | 91.8 | **95.7** | **96.3** | **94.6** | 96.1 |
| USNM374837 | 79.7 | 91.2 | **94.9** | **95.6** | **95.7** | 22.8 |
| USNM357999 | 79.9 | 91.2 | **94.9** | **95.5** | **95.6** | 95.6 |
| AMIB4247 | 79.7 | 91.2 | **94.8** | **95.5** | **95.7** | 9.3 |
| AMI33715-016 | 79.7 | 91.2 | **94.9** | **95.5** | **95.6** | 13.7 |
| USNM203781 | 79.6 | 91.2 | **95** | **95.6** | **95.6** | 2.5 |
| CAS28515 | 79.6 | 91.1 | **94.9** | **95.5** | **95.6** | 5.6 |
| Stubifer_GRA.1 | 79.9 | 91.1 | **94.7** | **95.4** | **95.5** | 95.8 |
| CAS247233.2 | 79.9 | 91.1 | **94.7** | **95.4** | **95.5** | 96.4 |
| USNM357892 | 79.6 | 91 | **94.7** | **95.4** | **95.6** | 3.9 |
| Stubifer_M118 | 80 | 91 | **94.6** | **95.3** | **95.3** | 96.2 |
| CAS225045 | 80.1 | 91 | **94.7** | **95.3** | **95.3** | 96.1 |
| AMI18353-041 | 79.4 | 91.1 | **94.8** | **95.5** | **95.5** | 2.6 |
| USNM349778 | 79.9 | 91 | **94.6** | **95.2** | **95.3** | 95.8 |
| CAS223855 | 79.4 | 91 | **94.7** | **95.3** | **95.5** | 6.8 |
| USNM396981 | 79.9 | 90.9 | **94.6** | **95.2** | **95.3** | 96.1 |
| USNM341594 | 79.8 | 90.9 | **94.6** | **95.2** | **95.3** | 61.4 |
| AMI19450-018.2 | 79.4 | 91.1 | **94.7** | **95.4** | **95.4** | 45.3 |
| USNM430718 | 79.8 | 91 | **94.5** | **95.2** | **95.3** | 95.3 |
| USNM357884 | 79.6 | 90.8 | **94.5** | **95.2** | **95.5** | 33.3 |
| CAS223939 | 79.8 | 91.3 | **95** | 93.2 | **95.8** | 12 |
| USNM357889 | 79.5 | 90.8 | **94.5** | **95.1** | **95.3** | 42.7 |
| USNM358001 | 79.3 | 90.6 | **94.5** | **95.1** | **95.3** | 3.2 |
| AMI20753-031 | 79.6 | 90.7 | 94.3 | **94.9** | **95.1** | 14.9 |
| USNM341595 | 79.8 | 90.5 | 94 | **94.6** | **94.7** | 95.5 |
| USNM374480 | 79.6 | 90.5 | 94 | **94.6** | **94.8** | 40.5 |
| AMI37933-007 | 79.5 | 90.2 | 94 | **94.6** | **95** | 1.9 |
| CAS223978 | 79.9 | 91 | **94.6** | 88.6 | **95.4** | 95.4 |
| CAS223979 | 78.9 | 89.5 | 93.3 | 93.9 | 94.2 | 17.7 |
| USNM298542 | 78.6 | 89.4 | 93.3 | 93.7 | 94 | 0.1 |
| USNM357897 | 79.4 | 89.4 | 93 | 93.5 | 93.7 | 95 |
| USNM112099 | 78 | 89.3 | 93.3 | 93.8 | 94.1 | 3.7 |
| AMI19450-018.1 | 78.1 | 89.4 | 93.1 | 93.6 | 94 | 2.6 |
| AMIB4208 | 80.7 | 90.3 | 94 | **94.6** | 90 | 6.7 |
| AMI20353-001 | 77.5 | 87.8 | 91.8 | 92.1 | 92.6 | 2.3 |
| AMI18740-066 | 80.4 | 83.3 | 86.8 | 84.8 | 85.4 | 2.1 |

Single nucleotide polymorphisms (SNPs) were detected for the light organ symbionts from most of the specimens sampled, but this number varied greatly and correlated with the variability in genome coverage (Spearman’s rank correlation: rho=0.84, P<0.001, Table S1). Samples with greater than 50% symbiont genome coverage at 10x read depth had an average of 23,221 SNPs relative to the reference genome of *P. mandapamensis* (Urbanczyk et al., 2011). A core set of 1,471 SNPs were identified across 32 specimens that represent 11 *Siphamia* host species and included reference genomes from both Clade I and Clade II of *Photobacterium mandapamensis.* 68% of these SNPs were synonymous, and the remaining non-synonymous SNPs were found in 288 distinct genes. None of the core SNPs were located in the *lux* operon, composed of the genes responsible for light production. However, two non-synonymous SNPs were detected in the *rpoN* gene, which is known to play a role in biofilm formation, bioluminescence, and symbiosis initiation for *Aliivibrio fischeri* (Wolfe et al., 2004), the luminous symbiont of many squid and other fish species. No other SNPs were detected in genes of known function for the bioluminescent symbiosis between *A. fischeri* and the squid host *Euprymna scolopes* (Norsworthy and Visick 2013).

A maximum likelihood phylogeny was inferred for the bacterial symbionts using full sequence alignments that included the core set of SNPs described above. This analysis confirmed that all *Siphamia* light organ symbionts examined belong to Clade II of *P. mandapamensis* and that the reference strain of *P. mandapamensis* representing Clade I (ajapo4.1) was a clear outgroup (Fig 4). The majority of symbionts analyzed were closely related to the reference strain svers1.1 of *P. mandapamensis,* although several symbionts fell out in a group with *P. mandapamensis* strain Res 4.1, both of which are members of Clade II. There were three additional symbionts, all from different host species, that did not belong to either of these subgroups, but are still clearly members of Clade II.

**Figure 4.**
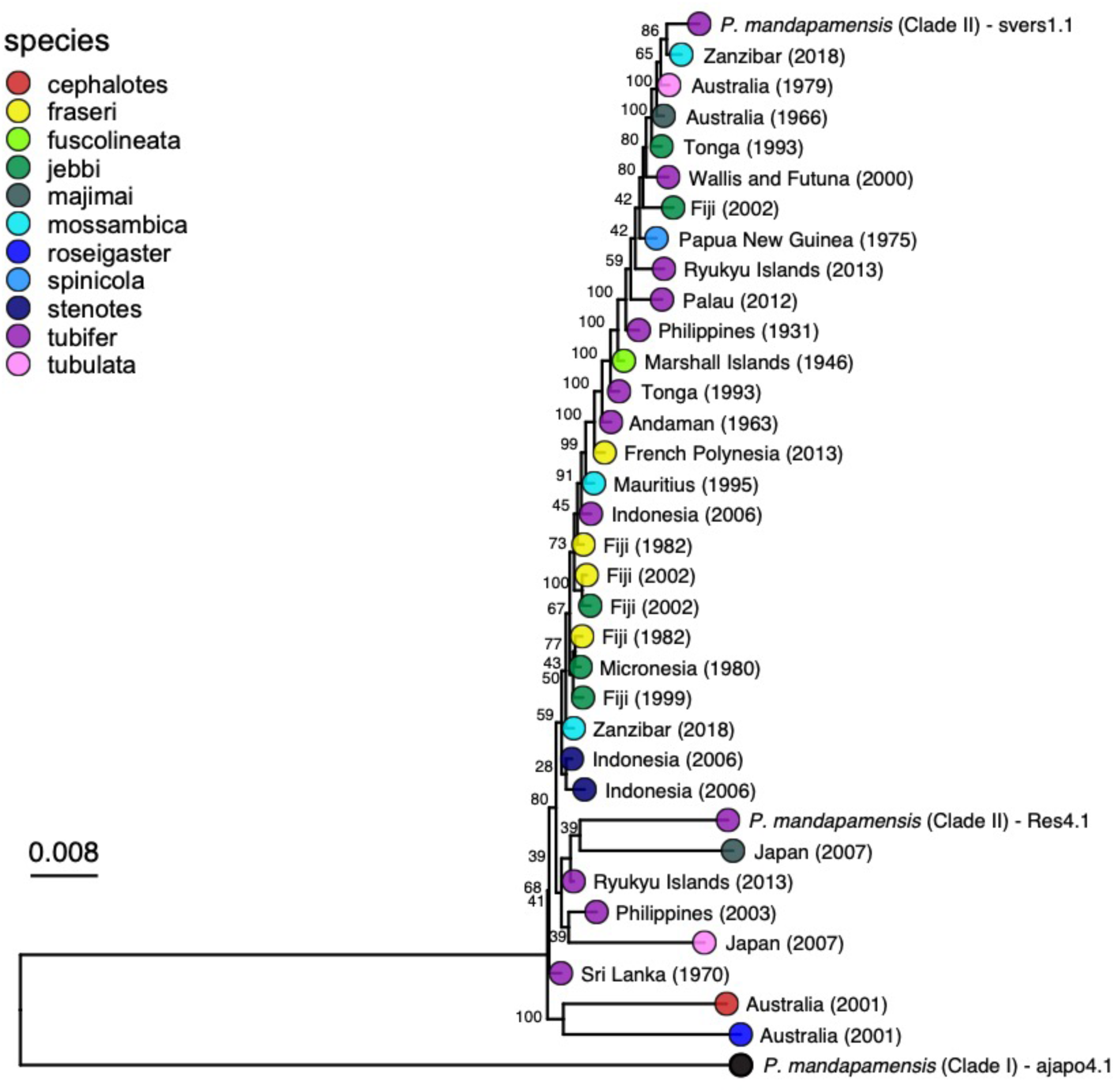
Maximum likelihood phylogeny of the light organ symbionts of various *Siphamia* species constructed from a core set of 1,471 single nucleotide polymorphisms. Corresponding host species are indicated by the branch tip colors and the sampling location and year of each specimen is listed in the branch label. Bootstrap support values are indicated at each node.

No clear patterns of symbiont divergence that corresponded with host species, geography, or time emerged. There was no correlation between symbiont phylogenetic distance and geographic distance (Spearman’s rank correlation: rho=-0.013, P_corr_,=1) and there was a slightly negative correlation between phylogenetic distance and time in years (Spearman’s rank correlation: rho=-0.17, P_corr_=0.006). In fact, the oldest specimen for which informative sequence data was retained was collected in 1931 and it had luminous bacteria in its light organ that was highly similar to symbionts from specimens collected more than eighty years later. Similarly, *Siphamia* specimens collected from locations in the western Indian Ocean had symbionts that were closely related to those from locations as far east as Fiji and even French Polynesia. With respect to S. *tubifer,* which has the broadest geographic distribution of all *Siphamia* species, the symbionts of all ten specimens included in the symbiont phylogeny fell out in Clade II of *P. mandapamensis* and showed no pattern of strain diversity by geography, confirming the high degree of specificity of this association, even across a broad geographic range.

The bacterial symbionts from four distinct host species had notably longer branches than the others, two of which were closely related to reference strain Res 4.1 (NZ_PYNS00000000), an isolate from the light organ of S. *tubifer* collected in Okinawa, Japan in 2014. Corresponding with longer branch lengths, these four symbionts had more than 3 times as many SNPs than any other sample, ranging between 66,583 and 72,219 SNPs (Table S1). Interestingly, these four specimens were collected from two locations, Sydney, Australia and Kochi, Japan, which had the lowest minimum annual temperatures of all collection sites in this study (Table 1).

Furthermore, there were 20,082 SNPs in common among these samples that were not present in the core set of SNPs identified across all samples.

### Analysis of co-divergence

Twenty specimens had informative sequence information for both the host and symbiont, and thus, we were able to carry out an analysis of co-divergence based on the host *COI* phylogeny and corresponding symbiont phylogeny for these individuals. This analysis revealed no evidence of co-divergence of *Siphamia* hosts and their light organ symbionts (P=0.13) as seen in Figure 5. However, S. *roseigaster* and S. *cephalotes* fall out as sister lineages relative to the rest of *Siphamia*, and their symbionts follow a similar pattern, forming a sister clade to the rest of *P. mandapamensis* Clade II.

**Figure 5.**
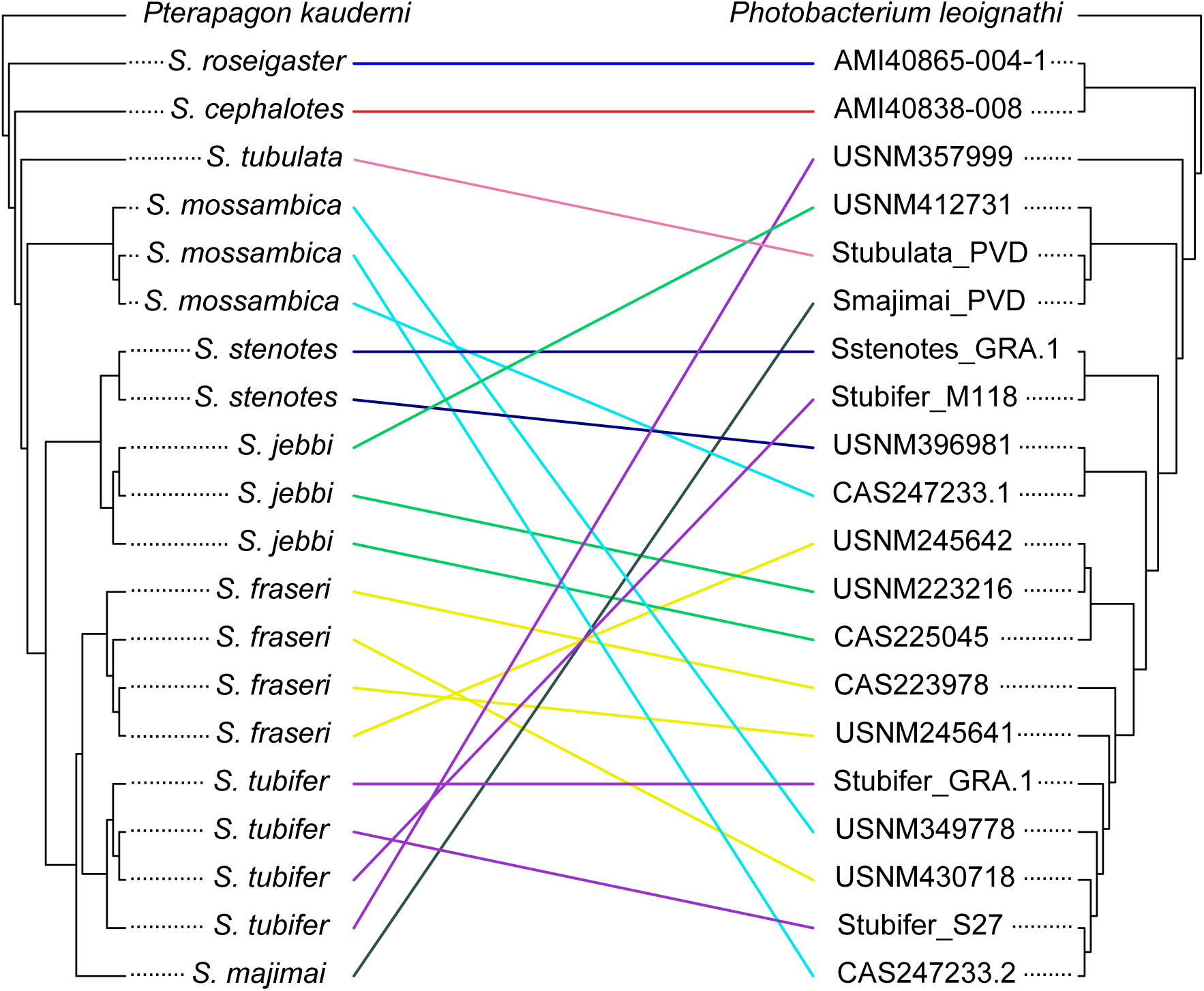
Analysis of the phylogenetic relationships of *Siphamia* hosts (left) and their light organ symbionts (right) revealed no evidence of co-divergence. The host cladogram is based on *COI* gene sequences and the symbiont cladogram is based on a core set of 1,471 single nucleotide polymorphisms. Linkages between individual hosts and their symbionts are shown and colored according to host species.

## Discussion

Our results indicate that the symbiosis between cardinalfishes in the genus *Siphamia* and the luminous bacterium *Photobacterium mandapamensis* is highly conserved across host species, over geographic space, and through time. All light organ symbionts examined were identified as strains belonging to Clade II of *P. mandapamensis* (Kaeding et al., 2007). This high subspecies level of specificity is surprising given the facultative symbiotic life history of the bacterium and the broad geographic and temporal ranges examined. Such a high degree of specificity is expected for vertically transmitted symbioses in which a host directly transfers its symbiotic bacteria to its offspring (Moran 2006). For environmentally transmitted symbioses where the specific association must be re-established by each new host generation, we expected a lower degree of specificity, similar to what has been documented for most other symbiotically luminous fishes such as the leiognathid fishes (Kaeding et al., 2007). Thus, the highly conserved relationship between *Siphamia* hosts and *P. mandapamensis* (Clade II) indicates there may be unique mechanisms in the host and/or symbiont that contribute to maintaining the specificity of the association.

*Siphamia tubifer* larvae only take up symbionts in the pelagic phase, when their light organ becomes receptive to colonization (Dunlap et al., 2012). Yet, *Photobacterium* spp. normally occurs in relatively low concentrations in the pelagic environment (Trousselier *et al.,* 2017), and even more so at the sub-species level. For the larval host to rely on this improbable encounter in the open water would be considered a very risky strategy. However, it has been shown that established populations of S. *tubifer* hosts regularly excrete their luminous symbiont with fecal waste (Dunlap & Nakamura 2011), thereby enriching its population in the immediate environment. Indeed, a previous study of S. *tubifer* symbiont genomics revealed fine-scale population structure of *P. mandapamensis* among geographic locations, indicating that symbiont populations are heavily influenced by their local hosts (Gould and Dunlap 2019). Therefore, this local enrichment may be a key mechanism/factor in mitigating the risk of relying on environmental transmission in the Siphamia-symbiont dependency, and for *Siphamia* hosts, ensures that *P. mandapamensis* (Clade II) will be readily available to new recruits anywhere that adult *Siphamia* already occur.

The apparent preference to associate with *P. mandapamensis* Clade II over strains in Clade I also suggests that there are critical strain level differences between members of these clades that may be of consequence to the host. However, most studies of microbial symbioses overlook symbiont strain-level variation, even though this variation can have important impacts on a host, and merits further investigation. For example, in *A. fischeri,* the primary symbiont for most *Euprymna* squid species, patterns of strain variation have been observed within and between host populations (Jones et al., 2006, Wollenberg and Ruby 2009), and can have different colonization efficiencies (Lee and Ruby 1994, Bongrand *et al.,* 2016), mechanisms of biofilm formation during host colonization (Rotman *et al.,* 2019), and may have variable fitness consequences to their host (Koch et al., 2014). In this study we were able to characterize strain variation in *P. mandapamensis* associated with various *Siphamia* hosts. There was no distinct correlation between symbiont strain and host species with respect to time or geography, although we did observe some strain divergence associated with colder temperatures. Four of the *Siphamia* specimens examined had more than three times as many symbiont SNPs as the others. Interestingly, these four individuals were all collected from more temperate regions in Japan and Australia with the lowest minimum annual temperatures of all locations in this study (Table 1). Temperature is a driving factor of the distribution of bacteria in the marine environment (Sul et al 2013), and has been shown to affect the distribution of the luminous vibrio symbionts of sepiolid squid (Nishiguchi 2000) and to regulate the symbiotic associations of other marine taxa, such as cnidarians (Herrera et al 2020). Thus, the symbionts associated with these four specimens might have some genetic adaptations to slightly cooler temperatures. Future studies investigating the influence of temperature on strain diversity and host colonization efficiency would help to elucidate the role that temperature might play in the *Siphamia-Photobacterium mandapamensis* symbiosis.

Our primary objective of this study was to sequence the symbionts found in the light organs of various *Siphamia* species, but we were able to recover enough host sequence data to also construct a reasonably well-supported host phylogeny. This allowed us to examine co­diversification of hosts and their microbial symbionts. Although we found no evidence of co­diversification, the high degree of specificity maintained for this symbiosis across host species through time and space suggests that this association is genetically constrained by the host. This host-mediated selection poses the question of whether *P. mandapamensis* (Clade II) provides a fitness advantage to the host compared to other bacteria moving through the gut of *Siphamia,* including other luminous bacteria. It should also be noted that a lack of co­diversification does not preclude a history of co-evolution of host and symbiont in the system (Moran 2006) and members of Clade II of *P. mandapamensis* are likely have specific adaptations that provide them with a fitness advantage inside the light organ environment of *Siphamia* fishes.

*Siphamia,* the only symbiotically luminous genus of cardinalfish, is monophyletic and divergent from the rest of the Apogonidae (Thacker and Roje 2009). The absence of this symbiosis in all other cardinalfish genera, including the other bioluminescent genera, brings up intriguing questions regarding the role of the symbiosis in the evolution of the Siphamia genus, specifically whether this association is a form of speciation by symbiosis (Wallin 1927), endowing *Siphamia* species with a key innovation that helped them persist and perhaps even proliferate. Parallel examples have been documented in damselfishes’ (Pomacentridae) mutualism with sea anemones, proposed to be the key innovation leading to the radiation of anemonefishes (Amphiprioninae; Litsios et al., 2012). Similarly, symbiosis with zooxanthellae may be a key attribute in enhancing adaptive radiation for the heterobranch genus *Phyllodesmium* (Wagele 2004). In *Siphamia,* there seems to be a rigorous mechanism of maintaining symbiont specificity across the host genus, presumably driven by the host. Therefore, understanding the genetic architecture of the Siphamia symbiont selection mechanism may be key to deciphering the highly specific nature of the association.

We also highlight the potential for formalin-fixed, fluid-preserved museum specimens to be used to study microbial symbioses. Adapting recently developed molecular techniques to extract and prepare DNA from these specimens for sequencing, including the use of single-stranded DNA as templates to construct sequence libraries, we recovered informative sequence data for both the host and its bacterial symbiont. This process allowed us to identify and compare strain level differences between the bacterial symbionts of many host species collected over nearly a century throughout the Indo-Pacific. We saw no clear correlation between sequence quality or yield and variables such as specimen age, size, or DNA input. It is likely that the observed variability between samples is largely due to the initial preservation method and long-term storage conditions of the specimens. For example, the quality (buffered or unbuffered) and concentration of the formalin solution used to initially fix a specimen can have variable effects on DNA quality (Hykin et al., 2015), as can the length of time a specimen remained in formalin before being transferred to its long-term storage solution. Unfortunately, many specimen records lack such information. Moving forward, it would be beneficial for researchers to have access to such information for specimens archived in natural history museums. Nevertheless, with the advancement of new genomic techniques and sequencing technologies, the ability to retrieve informative genetic information for both a host animal and its symbiotic bacteria from museum specimens will continue to advance our understanding of these critical associations.

## Data Availability Statement

Sequence data associated with this study will be made publicly available prior to publication.

## Author Contributions

ALG conceived of the project and secured funding for the work. AF-P and ALG carried out the genomic methods and analyses. AMG assisted with the identification of specimens and analysis of their associated metadata. All authors contributed to the discussion and interpretation of the results and to writing the manuscript. All authors approve of the submitted version of this manuscript.

## Acknowledgements

We would like to acknowledge Athena Lam and California Academy of Science’s Center for Comparative Genomics for technical support with our genomic methods as well as Joe Russack and James Henderson for their bioinformatics help and advice. Thank you to Luiz Rocha and Hudson Pinheiro for their collection efforts, Dave Catania and Mysi Hoang for their support with the museum specimens, and Jessica Herbert for her assistance in the lab. We also thank the Australian Museum and the Smithsonian National Museum of Natural History for access to their specimens for this project.

## Supplementary Information

**Figure S1.**
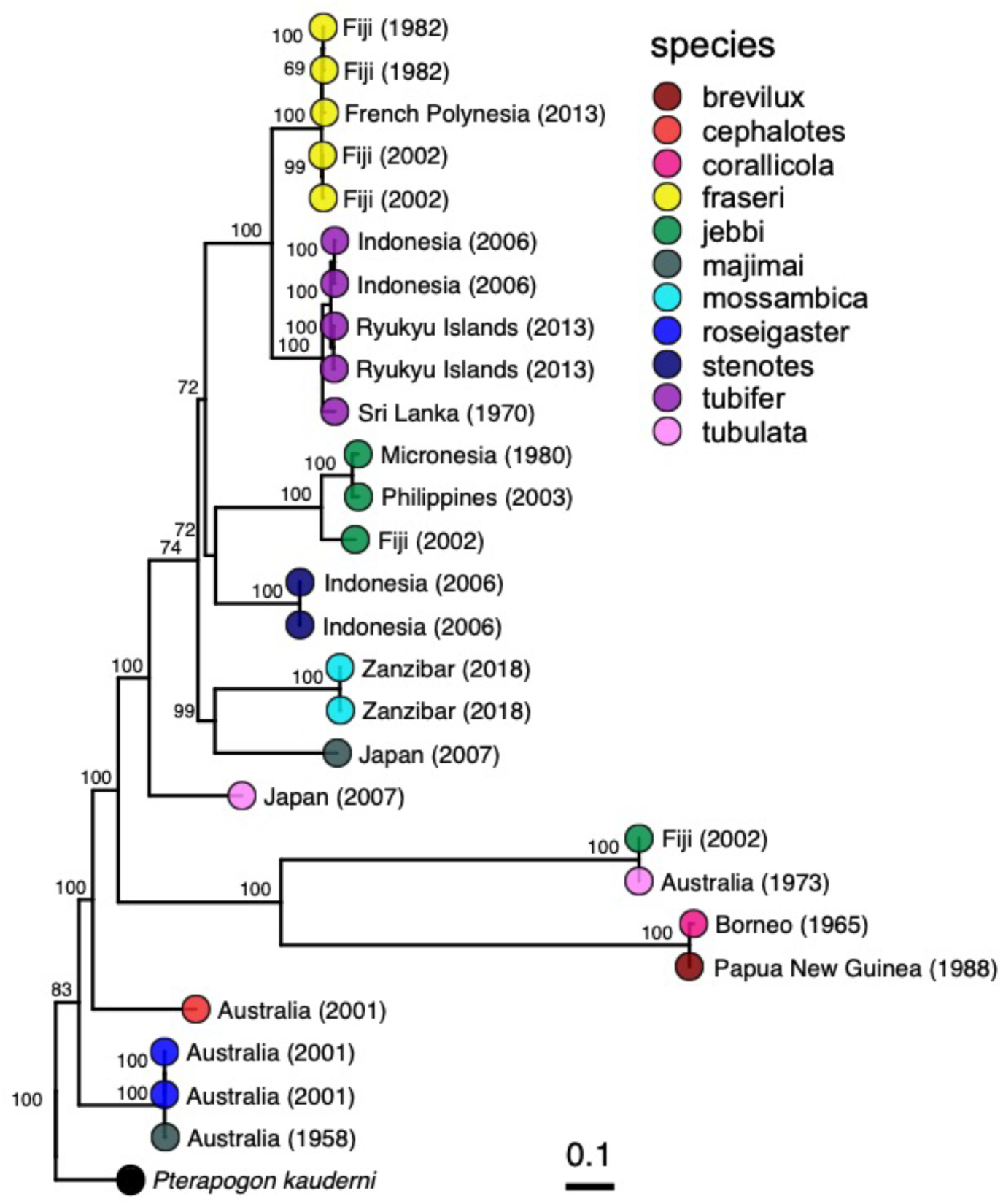
Maximum likelihood phylogeny of *Siphamia* based on a concatenated supermatrix of 15 mtDNA gene sequences *(ATP6, ATP8, COXI, COX2, COX3, CYTB, ND1, ND2, ND3, ND4, ND4L, ND5, ND6, 16S, 18S).* Species identities are indicated by the branch tip colors and the sampling location and year of each specimen is listed in the branch label.

**Table S1.** Information for the *Siphamia* specimens sampled and their corresponding sequence information. Listed are each specimen’s catalog number or unique identifier, species identification, sampling location and year, the standard length of the individual sampled, the total amount of double stranded DNA extracted from the light organ, the raw number of sequence reads, the number of reads that passed quality filtering and were trimmed, the number of reads that aligned to the symbiont reference genome *(P. mandapamensis* strain svers1.1), the percent of the symbiont reference genome covered at 10x sequence read depth, the total number of SNPs identified for each symbiont relative to the reference genome, the type of sequencing that was carried out, and the kit used for sequence library preparation. Specimens with decimals after their catalog number or unique identifier indicate that more than one individual was sampled from the same specimen lot.

| Specimen ID | Species | Location | Year | Length (cm) | Total dsDNA (ng) | Raw | Trimmed | Aligned | %10x | SNPs | Sequence Run | Library Prep |
| --- | --- | --- | --- | --- | --- | --- | --- | --- | --- | --- | --- | --- |
| AMI18353-041 | jebbi | Fiji | 1974 | 1.69 | <2 | 28258958 | 26861780 | 187425 | 0.5 | 0 | HiSeq 2x150 | Swift |
|  |  |  |  |  |  | 27257752 | 26799476 | 407326 | 1 | 166 | NovaSeq 2x150 | Swift |
| AMI18740-066 | jebbi | Australia | 1975 | 1.46 | <2 | 40906024 | 39209732 | 306641 | 2.1 | 0 | HiSeq 2x150 | Swift |
| AMI19450-018.1 | tubifer | Australia | 1975 | 2.94 | 3 | 58651761 | 56851489 | 442960 | 2.6 | 0 | HiSeq 2x150 | Swift |
| AMI19450-018.2 | tubifer | Australia | 1975 | 3.62 | 3.5 | 33895456 | 32610468 | 1337245 | 23.2 | 10 | HiSeq 2x150 | NEB Ultra II |
|  |  |  |  |  |  | 24526375 | 23858943 | 1042258 | 16.4 | 5,435 | NovaSeq 2x150 | NEB Ultra II |
| AMI20353-001 | majimai | Australia | 1972 | 1.67 | <2 | 39503975 | 38494612 | 344674 | 2.3 | 1 | HiSeq 2x150 | Swift |
| AMI20753-031 | tubulata | Australia | 1979 | 2.56 | <2 | 35965027 | 35217790 | 1085963 | 14.9 | 6,916 | NovaSeq 2x150 | Swift |
| AMI33715-016 | jebbi | Australia | 1993 | 1.46 | 2.2 | 35229540 | 33924547 | 359824 | 1.9 | 0 | HiSeq 2x150 | Swift |
|  |  |  |  |  |  | 33504802 | 33190608 | 655297 | 5 | 862 | NovaSeq 2x150 | Swift |
| AMI37933-007 | tubifer | Vanuatu | 1997 | 2.19 | 52.1 | 3604557 | 2090383 | 219645 | 0.1 | 47 | NovaSeq 2x150 | NEB Ultra II |
|  |  |  |  |  |  | 49624926 | 48314012 | 498902 | 1.9 | 0 | HiSeq 2x150 | Swift |
| AMI40838-008 | cephalotes | Australia | 2001 | 3.07 | 52.8 | 34771013 | 34092047 | 5099255 | 92.3 | 70,709 | NovaSeq 2x150 | NEB Ultra II |
| AMI40865-004.1 | roseigaster | Australia | 2001 | 4.56 | 74.1 | 91467705 | 89885499 | 6944692 | 93.4 | 72,219 | NovaSeq 2x150 | NEB Ultra II |
| AMI40865-004.2 | roseigaster | Australia | 2001 | 4.55 | 4.6 | 48426323 | 46531047 | 654416 | 5.9 | 1,588 | HiSeq 2x150 | Swift |
| AMIB4208 | majimai | Australia | 1958 | 2.31 | <2 | 36877472 | 34915450 | 467033 | 6.4 | 2 | HiSeq 2x150 | Swift |
|  |  |  |  |  |  | 87751 | 55995 | 12134 | 0 | 0 | NovaSeq 2x150 | Swift |
| AMIB4247 | tubifer | Vanuatu | 1959 | 2.01 | <2 | 28358555 | 27422784 | 269641 | 2 | 0 | HiSeq 2x150 | Swift |
|  |  |  |  |  |  | 23520680 | 23130431 | 584903 | 2.9 | 1,633 | NovaSeq 2x150 | Swift |
| CAS247233.1 | mossambica | Zanzibar | 2018 | 2.55 | 542 | 5478617 | 5384608 | 454932 | 76.9 | 8,112 | HiSeq 1x150 | SparQ |
| CAS247233.2 | mossambica | Zanzibar | 2018 | 2.92 | 1530 | 75836474 | 73645413 | 33775734 | 96.4 | 15,579 | NovaSeq 2x150 | NEB Ultra II |
| CAS222309 | jebbi | Fiji | 2002 | - | 10.2 | 50665038 | 45699796 | 28523692 | 95.2 | 21,238 | NovaSeq 2x150 | NEB Ultra II |
| CAS223855 | jebbi | Fiji | 2002 | - | <2 | 44113541 | 41855479 | 377420 | 2.4 | 3 | HiSeq 2x150 | Swift |
|  |  |  |  |  |  | 33488272 | 33006205 | 576345 | 2.8 | 469 | NovaSeq 2x150 | Swift |
| CAS223939.1 | jebbi | Fiji | 2002 | 2.35 | 11.8 | 28012894 | 21498132 | 225577 | 4.3 | 63 | HiSeq 1x150 | SparQ |
| CAS223939.2 | jebbi | Fiji | 2002 | 1.81 | 8.9 | 11046726 | 10551137 | 716650 | 12 | 7,698 | NovaSeq 2x150 | NEB Ultra II |
| CAS223978.1 | unknown | Fiji | 2002 | 3.68 | 178.5 | 330421 | 323250 | 32364 | 0.1 | 10 | HiSeq 1x150 | SparQ |
| CAS223978.2 | unknown | Fiji | 2002 | 4.05 | 50.5 | 23063434 | 21244678 | 10747171 | 95.4 | 18,773 | NovaSeq 2x150 | NEB Ultra II |
| CAS223979.1 | fraseri | Fiji | 2002 | 2.8 | 9.2 | 24192691 | 22748353 | 328578 | 0.9 | 0 | HiSeq 2x150 | NEB Ultra II |
|  |  |  |  |  |  | 26735318 | 25635641 | 457797 | 1.1 | 7 | NovaSeq 2x150 | NEB Ultra II |
| CAS223979.2 | fraseri | Fiji | 2002 | 3.04 | 15.5 | 10467851 | 9981063 | 381078 | 17.7 | 72 | HiSeq 1x150 | SparQ |
| CAS225045 | jebbi | Fiji | 1999 |  | 3.4 | 21590245 | 20304117 | 15184111 | 96.1 | 18,316 | NovaSeq 2x150 | NEB Ultra II |
| CAS27441 | tubifer | Philippines | 1931 | 3.26 | 1.8 | 18607665 | 18124484 | 1483134 | 95.3 | 18,841 | HiSeq 1x150 | SparQ |
| CAS28515 | tubulata | Australia | 1973 | - | <2 | 36559960 | 33708337 | 213670 | 1 | 2 | HiSeq 2x150 | Swift |
|  |  |  |  |  |  | 35922973 | 35323462 | 589696 | 3.1 | 1,301 | NovaSeq 2x150 | Swift |
| CAS84356 | tubifer | Palau | 2012 | 1.9 | 38.9 | 48287910 | 38767642 | 6258850 | 64.8 | 13,661 | HiSeq 1x150 | SparQ |
| Stubifer_M118 | tubifer | Ryukyu Islands | 2013 | 1.3 | 26.5 | 75237177 | 72661256 | 57068084 | 96.2 | 16,828 | HiSeq 2x150 | Swift |
| Smajimai_PVD | majimai | Japan | 2007 | 2.61 | 8949 | 36930530 | 35957911 | 26787137 | 94.8 | 70,889 | NovaSeq 2x150 | NEB Ultra II |
| Stubulata_PVD | tubulata | Japan | 2007 | 2.12 | 1225.5 | 34195315 | 33077310 | 21855337 | 95.1 | 66,583 | NovaSeq 2x150 | NEB Ultra II |
| Stubifer_S27 | tubifer | Ryukyu Islands | 2013 | 2.65 | - | 7403111 | 7277173 | 517331 | 88.9 | 19,790 | HiSeq 1x150 | SparQ |
| Sstenotes_GRA.1 | stenotes | Indonesia | 2006 | 1.89 | 115.9 | 22315393 | 24640343 | 21426979 | 95.8 | 24,716 | NovaSeq 2x150 | NEB Ultra II |
| Sstenotes_GRA.2 | stenotes | Indonesia | 2006 | 1.98 | 308 | 467708 | 426870 | 85208 | 0.5 | 23 | HiSeq 1x150 | SparQ |
| Stubifer_GRA.1 | tubifer | Indonesia | 2006 | 2.39 | 96 | 23365446 | 22298866 | 6159687 | 95.8 | 18,701 | NovaSeq 2x150 | NEB Ultra II |
| Stubifer_GRA.2 | tubifer | Indonesia | 2006 | 2.85 | 95.9 | 9455319 | 8509771 | 452475 | 52 | 3,041 | HiSeq 1x150 | SparQ |
| USNM112099 | elongata | Philippines | 1909 | 3.46 | <2 | 49945895 | 48231023 | 430697 | 3.7 | 8 | HiSeq 2x150 | Swift |
| USNM142281.1 | fuscolineata | Marshall Islands | 1946 | 2.2 | <2 | 16213148 | 15893522 | 6118510 | 96.1 | 15,366 | NovaSeq 2x150 | Swift |
| USNM142281.2 | fuscolineata | Marshall Islands | 1946 | 2.76 | 10.1 | 39670892 | 27192174 | 695872 | 62.8 | 381 | HiSeq 1x150 | NEB Ultra II |
| USNM203781 | corallicola | Borneo | 1965 | 2.58 | <2 | 33588104 | 32488841 | 640895 | 0.6 | 0 | HiSeq 2x150 | Swift |
|  |  |  |  |  |  | 56446868 | 51658133 | 1237586 | 1.2 | 178 | NovaSeq 2x150 | Swift |
| USNM223216 | jebbi | Micronesia | 1980 | 1.74 | 7 | 181380283 | 186040521 | 176746273 | 96.5 | 14,693 | NovaSeq 2x150 | NEB Ultra II |
| USNM245638 | jebbi | Fiji | 1982 | 2.07 | 2.2 | 36426337 | 34932628 | 508273 | 9.2 | 38 | HiSeq 2x150 | Swift |
|  |  |  |  |  |  | 27073152 | 26801480 | 647893 | 13 | 2,188 | NovaSeq 2x150 | Swift |
| USNM245641 | fraseri | Fiji | 1982 | 4.13 | 5.3 | 864701674 | 829515210 | 714629012 | 97.5 | 17,070* | NovaSeq 2x150 | NEB Ultra II |
| USNM245642 | fraseri | Fiji | 1982 | 3.65 | 13 | 35743393 | 35774905 | 33716527 | 96.2 | 16,396 | NovaSeq 2x150 | NEB Ultra II |
| USNM298542 | brevilux | Papua New Guinea | 1988 | 2.24 | 21.1 | 662337 | 482006 | 148333 | 0.1 | 23 | NovaSeq 2x150 | NEB Ultra II |
|  |  |  |  |  |  | 49597285 | 47386221 | 486582 | 2.4 | 0 | HiSeq 2x150 | Swift |
| USNM341594 | jebbi | Tonga | 1993 | 1.91 | <2 | 27318206 | 27030306 | 1313295 | 61.4 | 7,995 | NovaSeq 2x150 | Swift |
| USNM341595 | tubifer | Tonga | 1993 | 3.87 | 7.8 | 18908174 | 17504661 | 4858215 | 94.2 | 725 | HiSeq 2x150 | NEB Ultra II |
|  |  |  |  |  |  | 11846770 | 11271717 | 2974122 | 88.1 | 17,900 | NovaSeq 2x150 | NEB Ultra II |
| USNM349778 | mossambica | Mauritius | 1995 | 2.36 | 15 | 33471029 | 32308812 | 12943283 | 95.8 | 18,219 | NovaSeq 2x150 | NEB Ultra II |
| USNM357884 | tubifer | Philippines | 1980 | 3.68 | 7.3 | 18374904 | 17191172 | 759889 | 8.5 | 1 | HiSeq 2x150 | NEB Ultra II |
|  |  |  |  |  |  | 13974418 | 12217495 | 879577 | 11.9 | 4,148 | NovaSeq 2x150 | NEB Ultra II |
| USNM357889 | spinicola | Papua New Guinea | 1975 | 3.11 | 4.1 | 21853900 | 21329666 | 2042137 | 42.7 | 12,366 | NovaSeq 2x150 | NEB Ultra II |
|  |  |  |  |  |  | 46425 | 42876 | 22 | 0 | 0 | HiSeq 2x150 | NEB Ultra II |
| USNM357892 | tubifer | Red Sea | 1969 | 3.35 | <2 | 35509177 | 33451543 | 215301 | 0.7 | 1 | HiSeq 2x150 | Swift |
|  |  |  |  |  |  | 27788827 | 27434566 | 542781 | 1.7 | 1,107 | NovaSeq 2x150 | Swift |
| USNM357897 | tubifer | Andaman | 1963 | 4.09 | 3.9 | 26178035 | 25619280 | 9568200 | 95 | 17,295 | NovaSeq 2x150 | NEB Ultra II |
| USNM357999 | tubifer | Sri Lanka | 1970 | 2.94 | <2 | 54251311 | 53731832 | 26374339 | 95.6 | 22,417 | NovaSeq 2x150 | Swift |
| USNM358001 | majimai | Philippines | 1978 | 2.1 | <2 | 38324209 | 36313447 | 254445 | 0.8 | 2 | HiSeq 2x150 | Swift |
|  |  |  |  |  |  | 33631537 | 33293817 | 428773 | 1.1 | 151 | NovaSeq 2x150 | Swift |
| USNM374480 | majimai | Australia | 1966 | 1.97 | 2.1 | 63381524 | 62370257 | 1400945 | 40.5 | 8,157 | NovaSeq 2x150 | Swift |
| USNM374837 | unknown | Wallis and Futuna | 2000 | 1.96 | 12.1 | 10549216 | 7951371 | 985084 | 22.8 | 9,465 | NovaSeq 2x150 | NEB Ultra II |
|  |  |  |  |  |  | 41856128 | 40302628 | 595029 | 10.7 | 11 | HiSeq 2x150 | Swift |
| USNM396981 | stenotes | Indonesia | 2006 | 1.89 | 153.9 | 36161679 | 35712980 | 34096922 | 96.1 | 16,892 | NovaSeq 2x150 | NEB Ultra II |
| USNM412731 | jebbi | Philippines | 2003 | 1.73 | 23.6 | 27071135 | 25964385 | 16604538 | 95.9 | 32,687 | NovaSeq 2x150 | NEB Ultra II |
| USNM430718 | fraseri | French Polynesia | 2013 | 3.34 | 58.9 | 18468382 | 17513981 | 2806955 | 95.3 | 20,592 | NovaSeq 2x150 | NEB Ultra II |

**Table S2.**
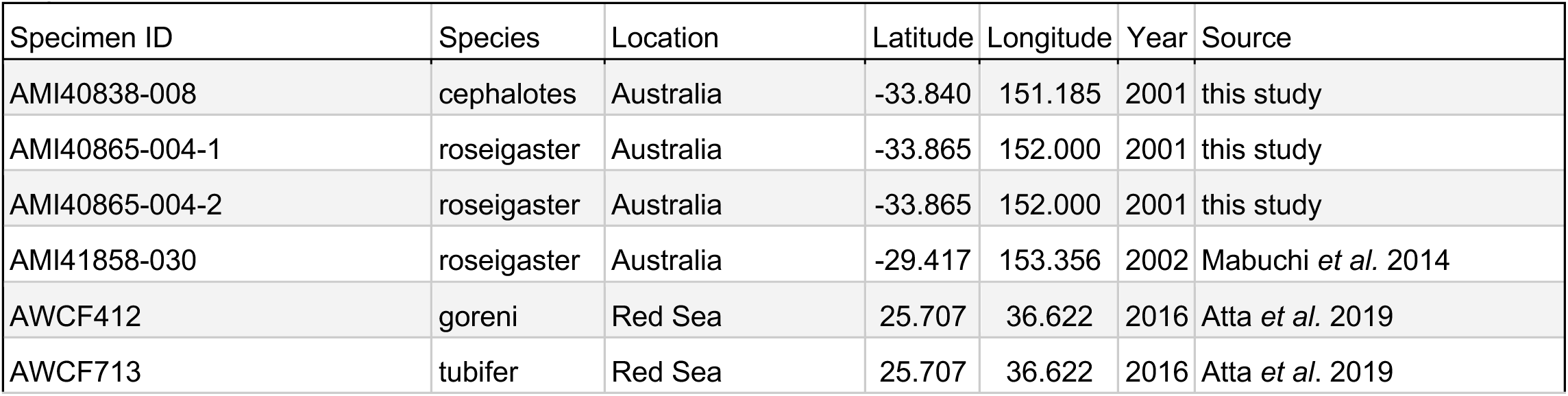
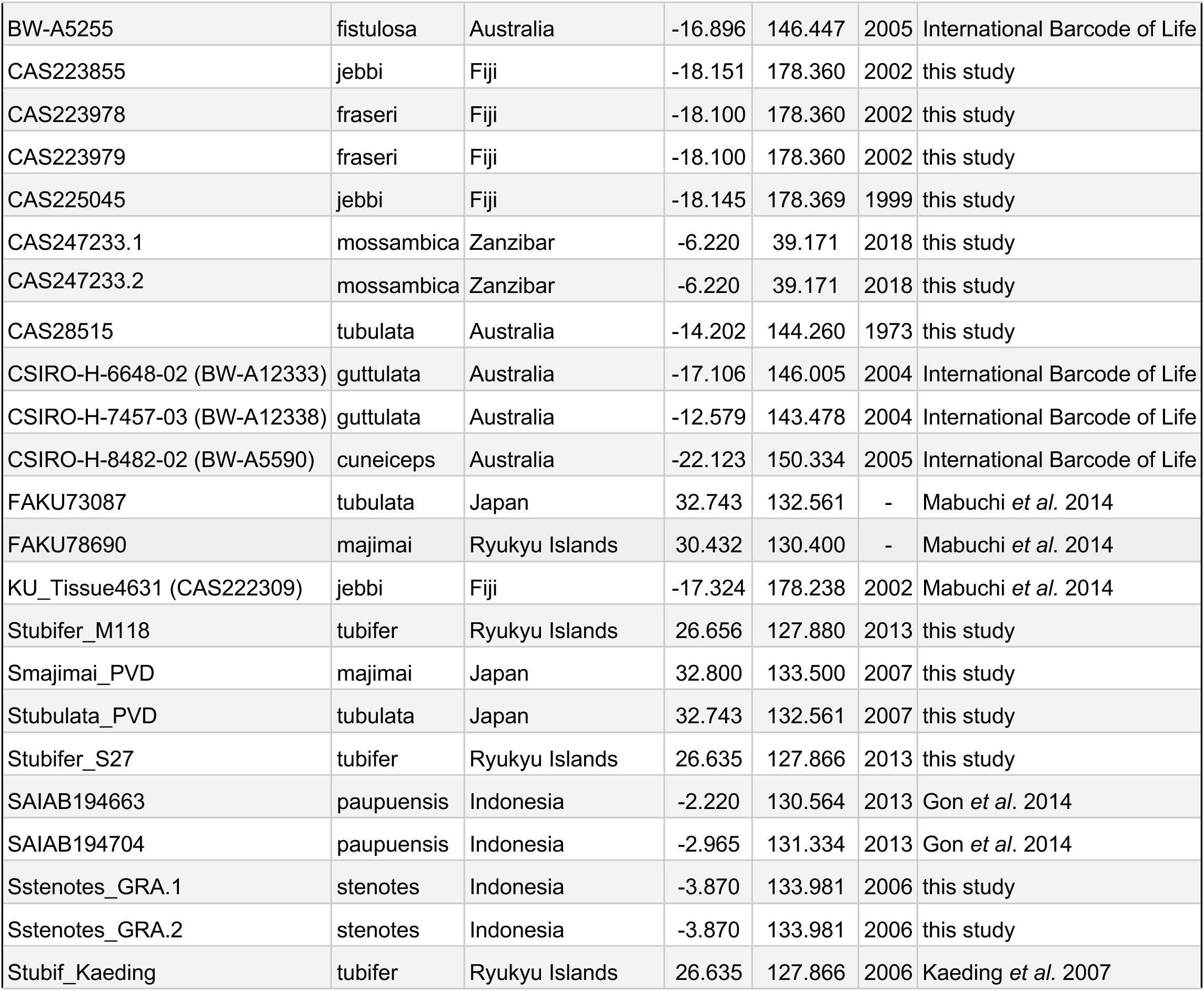
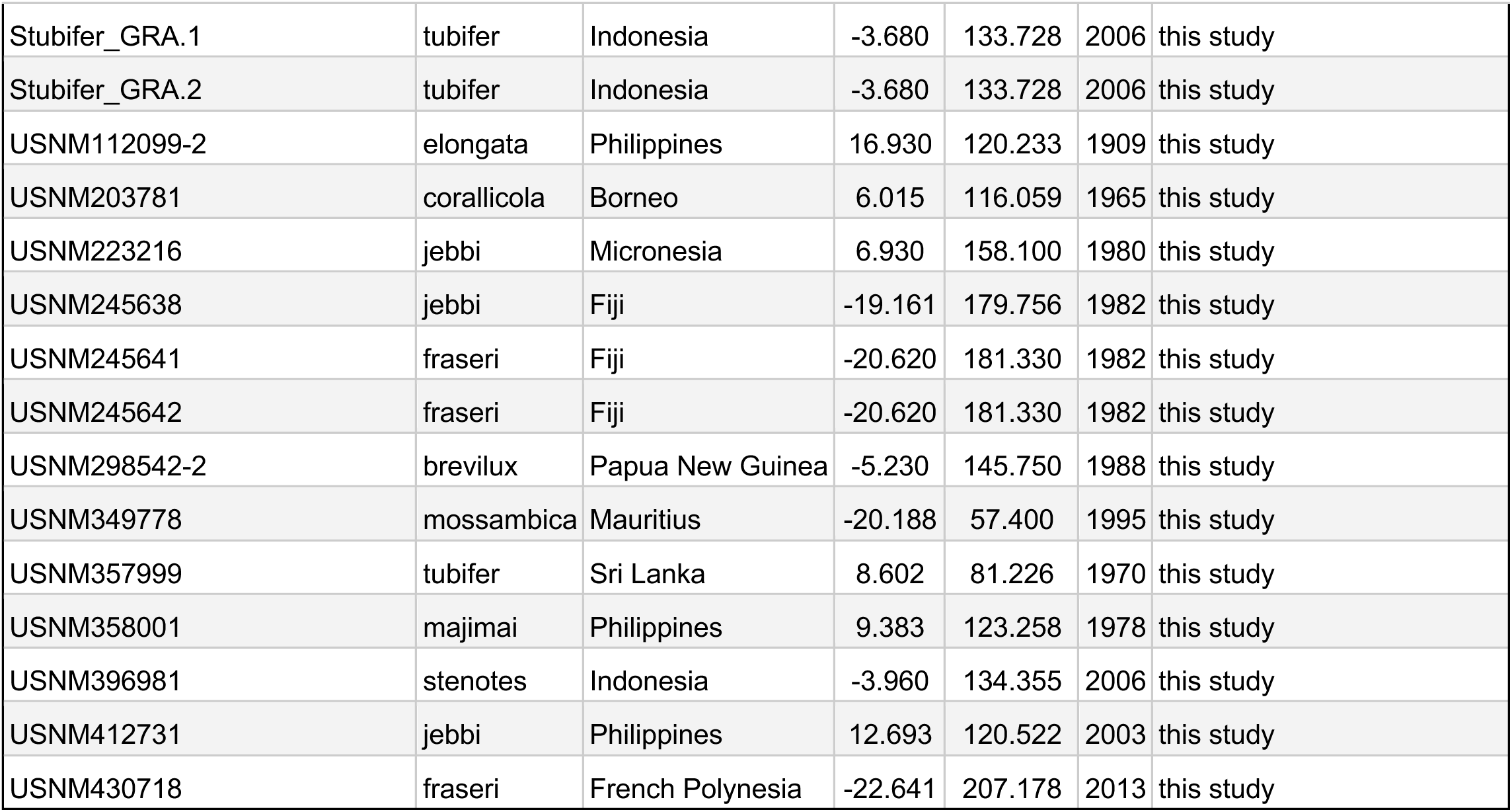
Information for the *Siphamia COI* sequences that were used to construct the host phylogeny. Listed are each specimen’s catalog number or unique identifier, species identification, sampling location, exact latitude and longitude, year, and the source of the sequence.

| Specimen ID | Species | Location | Latitude | Longitude | Year | Source |
| --- | --- | --- | --- | --- | --- | --- |
| AMI40838-008 | cephalotes | Australia | -33.840 | 151.185 | 2001 | this study |
| AMI40865-004-1 | roseigaster | Australia | -33.865 | 152.000 | 2001 | this study |
| AMI40865-004-2 | roseigaster | Australia | -33.865 | 152.000 | 2001 | this study |
| AMI41858-030 | roseigaster | Australia | -29.417 | 153.356 | 2002 | Mabuchi *et al.* 2014 |
| AWCF412 | goreni | Red Sea | 25.707 | 36.622 | 2016 | Atta *et al.* 2019 |
| AWCF713 | tubifer | Red Sea | 25.707 | 36.622 | 2016 | Atta *et al.* 2019 |
| BW-A5255 | fistulosa | Australia | -16.896 | 146.447 | 2005 | International Barcode of Life |
| CAS223855 | jebbi | Fiji | -18.151 | 178.360 | 2002 | this study |
| CAS223978 | fraseri | Fiji | -18.100 | 178.360 | 2002 | this study |
| CAS223979 | fraseri | Fiji | -18.100 | 178.360 | 2002 | this study |
| CAS225045 | jebbi | Fiji | -18.145 | 178.369 | 1999 | this study |
| CAS247233.1 | mossambica | Zanzibar | -6.220 | 39.171 | 2018 | this study |
| CAS247233.2 | mossambica | Zanzibar | -6.220 | 39.171 | 2018 | this study |
| CAS28515 | tubulata | Australia | -14.202 | 144.260 | 1973 | this study |
| CSIRO-H-6648-02 (BW-A12333) | guttulata | Australia | -17.106 | 146.005 | 2004 | International Barcode of Life |
| CSIRO-H-7457-03 (BW-A12338) | guttulata | Australia | -12.579 | 143.478 | 2004 | International Barcode of Life |
| CSIRO-H-8482-02 (BW-A5590) | cuneiceps | Australia | -22.123 | 150.334 | 2005 | International Barcode of Life |
| FAKU73087 | tubulata | Japan | 32.743 | 132.561 | - | Mabuchi *et al.* 2014 |
| FAKU78690 | majimai | Ryukyu Islands | 30.432 | 130.400 | - | Mabuchi *et al.* 2014 |
| KU_Tissue4631 (CAS222309) | jebbi | Fiji | -17.324 | 178.238 | 2002 | Mabuchi *et al.* 2014 |
| Stubifer_M118 | tubifer | Ryukyu Islands | 26.656 | 127.880 | 2013 | this study |
| Smajimai_PVD | majimai | Japan | 32.800 | 133.500 | 2007 | this study |
| Stubulata_PVD | tubulata | Japan | 32.743 | 132.561 | 2007 | this study |
| Stubifer_S27 | tubifer | Ryukyu Islands | 26.635 | 127.866 | 2013 | this study |
| SAIAB194663 | paupuensis | Indonesia | -2.220 | 130.564 | 2013 | Gon *et al.* 2014 |
| SAIAB194704 | paupuensis | Indonesia | -2.965 | 131.334 | 2013 | Gon *et al.* 2014 |
| Sstenotes_GRA.1 | stenotes | Indonesia | -3.870 | 133.981 | 2006 | this study |
| Sstenotes_GRA.2 | stenotes | Indonesia | -3.870 | 133.981 | 2006 | this study |
| Stubif_Kaeding | tubifer | Ryukyu Islands | 26.635 | 127.866 | 2006 | Kaeding *et al.* 2007 |
| Stubifer_GRA.1 | tubifer | Indonesia | -3.680 | 133.728 | 2006 | this study |
| Stubifer_GRA.2 | tubifer | Indonesia | -3.680 | 133.728 | 2006 | this study |
| USNM112099-2 | elongata | Philippines | 16.930 | 120.233 | 1909 | this study |
| USNM203781 | corallicola | Borneo | 6.015 | 116.059 | 1965 | this study |
| USNM223216 | jebbi | Micronesia | 6.930 | 158.100 | 1980 | this study |
| USNM245638 | jebbi | Fiji | -19.161 | 179.756 | 1982 | this study |
| USNM245641 | fraseri | Fiji | -20.620 | 181.330 | 1982 | this study |
| USNM245642 | fraseri | Fiji | -20.620 | 181.330 | 1982 | this study |
| USNM298542-2 | brevilux | Papua New Guinea | -5.230 | 145.750 | 1988 | this study |
| USNM349778 | mossambica | Mauritius | -20.188 | 57.400 | 1995 | this study |
| USNM357999 | tubifer | Sri Lanka | 8.602 | 81.226 | 1970 | this study |
| USNM358001 | majimai | Philippines | 9.383 | 123.258 | 1978 | this study |
| USNM396981 | stenotes | Indonesia | -3.960 | 134.355 | 2006 | this study |
| USNM412731 | jebbi | Philippines | 12.693 | 120.522 | 2003 | this study |
| USNM430718 | fraseri | French Polynesia | -22.641 | 207.178 | 2013 | this study |

**Table S3.** Results of the nucleotide BLAST search of symbiont 16S rRNA genes. Listed are each specimen’s catalog number or unique identifier, the percent of the reference 16S rRNA gene sequence *(Photobacterium leiognathi,* AY292917) covered at 10x sequence depth, the top matching sequence from the NCBI database including its accession number in parentheses, and the corresponding query coverage, E-value, and percent identity relative to that sequence.

| Specimen ID | %10x | Top hit | Query coverage | E-value | % identity |
| --- | --- | --- | --- | --- | --- |
| AMI18353-041 | 88.49 | Photobacterium leiognathi subsp. mandapamensis strain MahLm3 (JN380344.1) | 91% | 0.0 | 86.08 |
| AMI18740-066 | 89.2 | Photobacterium leiognathi (AY292917.1) | 93% | 0.0 | 79.97 |
| AMI19450-018.1 | 98.25 | Photobacterium leiognathi strain LC1-277 (AB243248.1) | 90% | 0.0 | 86.48 |
| AMI19450-018.2 | 99.94 | Photobacterium leiognathi strain ljone1.1 (AY204494.1) | 94% | 0.0 | 95.96 |
| AMI20353-001 | 95.02 | Photobacterium leiognathi strain W214 (MF554624.1) | 89% | 0.0 | 83.82 |
| AMI20753-031 | 99.94 | Photobacterium leiognathi strain ljone1.1 (AY204494.1) | 94% | 0.0 | 95.14 |
| AMI33715-016 | 100 | Photobacterium mandapamensis seaf.1.4 (AY455873.1) | 95% | 0.0 | 84.58 |
| AMI37933-007 | 99.81 | Photobacterium mandapamensis seaf.1.4 (AY455873.1) | 95% | 0.0 | 95.93 |
| AMI40838-008 | 100 | Photobacterium leiognathi subsp. mandapamensis strain ATCC 27561 (NR_115206.1) | 95% | 0.0 | 99.93 |
| AMI40865-004.1 | 99.94 | Photobacterium leiognathi subsp. mandapamensis strain ATCC 27561 (NR_115206.1) | 95% | 0.0 | 99.86 |
| AMI40865-004.2 | 99.94 | Photobacterium leiognathi strain AK-MIE (MH746214.1) | 91% | 0.0 | 99.01 |
| AMIB4208 | 100 | Photobacterium leiognathi strain LC1-283 (AB243249.1) | 90% | 0.0 | 90.74 |
| AMIB4247 | 99.94 | Photobacterium leiognathi strain LC1-283 (AB243249.1) | 90% | 0.0 | 92.75 |
| CAS222309 | 100 | Photobacterium leiognathi subsp. mandapamensis strain ATCC 27561 (NR_115206.1) | 95% | 0.0 | 100.00 |
| CAS223855 | 100 | Photobacterium leiognathi strain LC1-283 (AB243249.1) | 88% | 0.0 | 86.35 |
| CAS223939.1 | 99.94 | Photobacterium leiognathi strain ljone1.1 (AY204494.1) | 94% | 0.0 | 99.04 |
| CAS223939.2 | 99.94 | Photobacterium mandapamensis seaf.1.1 (AY455871.1) | 94% | 0.0 | 97.67 |
| CAS223978.1 | 29.61 | Photobacterium leiognathi subsp. mandapamensis strain ATCC 27561 (NR_115206.1) | 95% | 0.0 | 99.73 |
| CAS223978.2 | 100 | Photobacterium mandapamensis seaf.1.1 (AY455871.1) | 94% | 0.0 | 100.00 |
| CAS223979.1 | 99.94 | Photobacterium leiognathi strain LC1-277 (AB243248.1) | 90% | 0.0 | 94.88 |
| CAS223979.2 | 99.94 | Photobacterium leiognathi subsp. mandapamensis strain ATCC 27561 (NR_115206.1) | 95% | 0.0 | 100.00 |
| CAS225045 | 100 | Photobacterium leiognathi subsp. mandapamensis strain ATCC 27561 (NR_115206.1) | 95% | 0.0 | 100.00 |
| CAS247233.1 | 100 | Photobacterium mandapamensis seaf.1.1 (AY455871.1) | 94% | 0.0 | 100.00 |
| CAS247233.2 | 99.94 | Photobacterium leiognathi subsp. mandapamensis strain ATCC 27561 (NR_115206.1) | 95% | 0.0 | 100.00 |
| CAS27441 | 100 | Photobacterium leiognathi subsp. mandapamensis strain ATCC 27561 (NR_115206.1) | 95% | 0.0 | 100.00 |
| CAS28515 | 100 | Photobacterium leiognathi subsp. mandapamensis strain MahLm3 (JN380344.1) | 92% | 0.0 | 84.13 |
| CAS84356 | 99.94 | Photobacterium leiognathi subsp. mandapamensis strain ATCC 27561 (NR_115206.1) | 95% | 0.0 | 99.93 |
| Smajimai_PVD | 100 | Photobacterium leiognathi subsp. mandapamensis strain ATCC 27561 (NR_115206.1) | 95% | 0.0 | 99.93 |
| Sstenotes_GRA.1 | 100 | Photobacterium leiognathi subsp. mandapamensis strain ATCC 27561 (NR_115206.1) | 95% | 0.0 | 100.00 |
| Sstenotes_GRA.2 | 99.94 | Photobacterium leiognathi subsp. mandapamensis strain ATCC 27561 (NR_115206.1) | 95% | 0.0 | 99.73 |
| Stubifer_GRA.1 | 100 | Photobacterium leiognathi subsp. mandapamensis strain ATCC 27561 (NR_115206.1) | 95% | 0.0 | 100.00 |
| Stubifer_GRA.2 | 99.94 | Photobacterium leiognathi subsp. mandapamensis strain ATCC 27561 (NR_115206.1) | 95% | 0.0 | 99.93 |
| Stubifer_M118 | 100 | Photobacterium leiognathi subsp. mandapamensis strain ATCC 27561 (NR_115206.1) | 95% | 0.0 | 99.93 |
| Stubifer_S27 | 99.94 | Photobacterium leiognathi subsp. mandapamensis strain ATCC 27561 (NR_115206.1) | 95% | 0.0 | 100.00 |
| Stubulata_PVD | 100 | Photobacterium leiognathi subsp. mandapamensis strain ATCC 27561 (NR_115206.1) | 95% | 0.0 | 99.93 |
| USNM112099 | 92.18 | Photobacterium leiognathi strain AK5 (AB243232.1) | 90% | 0.0 | 85.57 |
| USNM142281.1 | 99.94 | Photobacterium leiognathi subsp. mandapamensis strain ATCC 27561 (NR_115206.1) | 95% | 0.0 | 100.00 |
| USNM142281.2 | 100 | Photobacterium leiognathi strain ljone1.1 (AY204494.1) | 93% | 0.0 | 94.36 |
| USNM203781 | 100 | Photobacterium leiognathi strain LC1-277 (AB243248.1) | 90% | 0.0 | 90.68 |
| USNM223216 | 100 | Photobacterium leiognathi subsp. mandapamensis strain ATCC 27561 (NR_115206.1) | 95% | 0.0 | 100.00 |
| USNM245638 | 100 | Photobacterium leiognathi strain AK5 (AB243232.1) | 90% | 0.0 | 93.11 |
| USNM245641 | 100 | Photobacterium leiognathi subsp. mandapamensis strain ATCC 27561 (NR_115206.1) | 95% | 0.0 | 99.93 |
| USNM245642 | 100 | Photobacterium leiognathi subsp. mandapamensis strain ATCC 27561 (NR_115206.1) | 95% | 0.0 | 100.00 |
| USNM298542 | 99.94 | Photobacterium leiognathi subsp. mandapamensis strain MahLm3 (JN380344.1) | 91% | 0.0 | 82.60 |
| USNM341594 | 100 | Photobacterium leiognathi subsp. mandapamensis strain MahLm3 (JN380344.1) | 92% | 0.0 | 93.40 |
| USNM341595 | 100 | Photobacterium leiognathi subsp. mandapamensis strain ATCC 27561 (NR_115206.1) | 95% | 0.0 | 100.00 |
| USNM349778 | 100 | Photobacterium leiognathi subsp. mandapamensis strain ATCC 27561 (NR_115206.1) | 95% | 0.0 | 99.93 |
| USNM357884 | 100 | Photobacterium mandapamensis se afl.1.1 (AY455871.1) | 94% | 0.0 | 100.00 |
| USNM357889 | 100 | Photobacterium mandapamensis se afl.1.4 (AY455873.1) | 95% | 0.0 | 97.90 |
| USNM357892 | 100 | Photobacterium leiognathi strain LC1-277 (AB243248.1) | 90% | 0.0 | 91.53 |
| USNM357897 | 100 | Photobacterium leiognathi subsp. mandapamensis strain ATCC 27561 (NR_115206.1) | 95% | 0.0 | 99.93 |
| USNM357999 | 100 | Photobacterium leiognathi subsp. mandapamensis strain ATCC 27561 (NR_115206.1) | 95% | 0.0 | 99.93 |
| USNM358001 | 100 | Photobacterium leiognathi strain LC1-277 (AB243248.1) | 90% | 0.0 | 87.62 |
| USNM374480 | 100 | Photobacterium leiognathi strain LC1-283 (AB243249.1) | 90% | 0.0 | 91.82 |
| USNM374837 | 100 | Photobacterium leiognathi subsp. mandapamensis strain MahLm3 (JN380344.1) | 91% | 0.0 | 87.07 |
| USNM396981 | 100 | Photobacterium leiognathi strain lleuc1.1 (AY204495.1) | 94% | 0.0 | 100.00 |
| USNM412731 | 100 | Photobacterium leiognathi subsp. mandapamensis strain ATCC 27561 (NR_115206.1) | 95% | 0.0 | 100.00 |
| USNM430718 | 99.94 | Photobacterium leiognathi subsp. mandapamensis strain ATCC 27561 (NR_115206.1) | 95% | 0.0 | 100.00 |

